# Arabidopsis epigenetic factor AS2 attenuates nucleolar stress by camptothecin and establishes leaf polarity by repressing a CDK inhibitor

**DOI:** 10.1101/2025.09.23.677680

**Authors:** Ayami Nakagawa, Hidekazu Iwakawa, Hiro Takahashi, Simon Vial-Pradel, Mari Takahashi, Kazuomi Ohga, Takuma Ito, Motoki Tamai, Eri Takada, Sumie Keta, Nanako Ishibashi, Sayuri Ando, Iwai Ohbayashi, Masaki Ito, Mami Yamazaki, Byung-Yoon Cha, Je-Tae Woo, Michiko Sasabe, Munetaka Sugiyama, Shoko Kojima, Yasunori Machida, Chiyoko Machida

**Affiliations:** Graduate School of Bioscience and Biotechnology, Chubu University, Kasugai 487-8501, Japan; Institute of Transformative Bio-Molecules (WPI-ITbM), Nagoya University, Nagoya 464-8602, Japan; Division of Biological Science and Technology, Graduate School of Natural Science and Technology, Kanazawa University Kakuma-machi, Kanazawa, 920-1192, Japan; Graduate School of Medical Sciences, Kanazawa University, Kakuma-machi, Kanazawa, Ishikawa 920-1192, Japan; Division of Biological Science, Graduate School of Science, Nagoya University, Nagoya 464-8602, Japan; Department of Life Sciences, National Cheng Kung University, Tainan City 701, Taiwan; Graduate School of Pharmaceutical Sciences, Chiba University, Chiba 260-8675, Japan; Department of Biology, Faculty of Agriculture and Life Science, Hirosaki University, Bunkyo-cho, Hirosaki 036-8561, Japan; Department of Biological Sciences, Graduate School of Science, The University of Tokyo, Tokyo 113-0033, Japan

**Keywords:** adaxial-abaxial leaf polarity, *ASYMMETRIC LEAVES2*, AS2 body, CDK inhibitor *KRP5*/*ICK3*, nucleolus, topoisomerase I

## Abstract

The *Arabidopsis thaliana* leaf, exhibiting a symmetrically extended flat morphology, consists of two distinct cellular domains: adaxial and abaxial layers. The *ASYMMETRIC LEAVES2* (*AS2*) gene is essential for specifying the adaxial domain, and its protein forms nucleolar structures, termed AS2 bodies, at ribosomal-DNA loci. Numerous nucleolus-related genes have been reported to be cooperatively involved in leaf adaxialization together with *AS2*. However, the molecular relationships between *AS2* and these genes remain unclear. To identify chemical modulators of AS2 function, we screened a chemical library containing natural products and identified eight molecules, including camptothecin, that induced filamentous leaves in the *as2* mutant. Camptothecin is an inhibitor of topoisomerase I, which is required for transcription of ribosomal–RNA genes, and induces nucleolar stress. Treatment with 10-hydroxy-camptothecin increased *KIP*-*RELATED*-*PROTEIN5*/*INHIBITOR*-*OF*-*CYCLIN*-*DEPENDENT*-*KINASE3* (*KRP5/ICK3*) transcript levels, encoding a CDK inhibitor, and caused notable changes in the number and morphology of AS2 bodies. Given its role in leaf morphogenesis, our findings suggest that AS2 is a key factor in establishing leaf adaxial-abaxial polarity by regulating cell proliferation and protecting against nucleolar stress through coordinated interactions with nucleolar proteins.

## INTRODUCTION

Morphogenesis of multi-cellular organisms tightly coordinates cell division and cell differentiation (Harashima and Schnittger, 2010). Leaves, lateral organs that emerge from a mass of stem cells (called a shoot apical meristem, SAM), grow and differentiate along three axes (proximal-distal, adaxial-abaxial, and medial-lateral) to form flat and symmetric leaves. Waites et al. identified the *PHANTASTICA* (*PHAN*) gene in *Antirrhinum majus* and suggested that the dorsal (adaxial) cell identity of leaves is required for lateral growth of the wild-type leaf (Waites and Hudson, 1995; Waites et al., 1998; Hudson, 2000). *ASYMMETRIC LEAVES1* (*AS1*) is ortholog of *PHAN* in *Arabidopsis thaliana* (Byrne et al., 2000). The initial leaf primordium is radial with abaxial cells, and some parts of the basal cells more adjacent to the top of the radially shaped SAM could initiate to differentiate into adaxially fated cells, establishing the leaf adaxial-abaxial polarity (Byrne et al., 2000; Iwakawa et al., 2002; Bowman 2002; Newman et al., 2002; Tsukaya 2006; Fukushima & Hasebe 2014). Then, after active division of cells locating along the medial-lateral axis, a flat and symmetric leaf is formed. Although the antagonistic gene interactions between the adaxial and abaxial polarity determinants should play a key role in generating leaf flatness (Wu et al., 2008; Iwasaki et al., 2013; Fukushima & Hasebe 2014; Machida et al., 2015; Husbands et al., 2015), it is still not known how these leaf polarity determinants regulate genes that control division of cells.

ASYMMETRIC LEAVES2 (AS2) protein (zinc-finger protein, a member of AS2/LOB family) forms a protein complex with the AS1 protein (Myb domain protein) (Byrne et al., 2000; Iwakawa et al., 2002; Guo et al. 2008; Yang et al. 2008) and they are colocalized to nucleoplasm, and form subnucleolar bodies called AS2 bodies at peripheral region of the nucleoli. (Ueno et al., 2007; Luo et al., 2012; 2020). AS2 bodies partially overlap with chromocenters containing nucleolus organizer regions (NORs), which consist of condensed and packed heterochromatin from 45S ribosomal DNA (rDNA) repeats (Luo et al., 2020; Iwakawa et al., 2020; Ando et al., 2023). The AS2-AS1 complex directly represses expression of the class 1 *KNOX* homeobox genes specifying the proximal-distal polarity of leaves in Arabidopsis (Yang et al., 2008; Guo et al., 2008; Ikezaki et al., 2010). The AS2-AS1 protein complex represses expression of the *ETTIN*/*AUXIN RESPONSE FACTOR3* (*ETT*/*ARF3*) gene and indirectly both *ETT*/*ARF3* and *ARF4* genes via function of miR390-tasiR-ARF (Iwasaki et al., 2013; Husbands et al., 2015). AS2-AS1 is involved in the maintenance of gene body DNA methylation in the *ETT*/*ARF3*. An inverse correlation has been shown between extents of levels of CpG methylation in the *ETT*/*ARF3* gene body and those of accumulation levels of its transcripts (Iwasaki et al., 2013; Vial-Pradel et al., 2018). Maintenance of repression of *ETT*/*ARF3* by AS2-AS1 is a crucial step for specification of the adaxial-abaxial polarity to form flat leaves (Iwasaki et al., 2013; Takahashi et al., 2013; Matsumura et al., 2016). Formation of the AS2 bodies at the periphery of the nucleolus is tightly related to the repression level of *ETT*/*ARF3* transcript (Luo et al., 2020; Ando et al., 2023).

Each of *as2* and *as1* single mutants exhibits a phenotype of weakly impaired leaf adaxialization (Iwakawa et al., 2007), additionally, when other gene mutations in the *as2* or *as1* background often exhibit filamentous (rod-shaped) leaves, which show complete loss of leaf adaxialization (Pinon et al., 2008; Yao et al., 2008; Szakonyi et al., 2011; Horiguchi et al., 2011; Iwasaki et al., 2013; Machida et al., 2015). Genetic screening with *as2* or *as1* mutants have identified 60 genes (hereinafter called ‘modifiers’) that are involved in leaf adaxialization with AS2-AS1 (Machida et al., 2015; Iwakawa et al., 2020). These modifier genes could be classified into several groups based on their function of encoded proteins; such as ribosome components, and nucleolar components, those involved in genome stability, chromatin structures, and tasiR-ARF biogenesis. Many of these modifier genes are related to the structural and functional components and functions of the nucleolus, such as genes encoding nucleolar proteins, *RNA HELICASE10* (*RH10*), *NUCLEOLIN1* (*NUC1*), *ROOT INITIATION DEFECTIVE2* (*RID2*), (Ohbayashi et al., 2011; 2017; Matsumura et al., 2016; Ohbayashi & Sugiyama 2018; Iwakawa et al., 2020). However, molecular relationship between AS2-AS1 which is involved in leaf development, and the modifier proteins remains unclear.

We performed chemical screening to clarify the relationship between the function of modifiers and leaf development involving AS2-AS1. Since many antibiotics for inhibiting function of ribosomal proteins are isolated from natural sources, and many naturally isolated molecules have been used as cancer suppressor drugs, which inhibit more active cell division (Boulon et al., 2010; Burger et al., 2013), we screened a chemical library containing natural products. As a result, eight molecules that induce filamentous leaf formation in the *as2* and *as1* mutants were isolated. Camptothecin (CPT) was identified as one of the molecules from the library. The application to plants of CPT, a well-known isoquinoline alkaloid developed as a specific inhibitor of topoisomerase I (TOPI) (Hsiang et al., 1985; 1989), might allow us to rapidly analyze leaf shapes and gene expression. It has also been reported that TOPI localized in the nucleolus is required for the transcription of ribosomal RNA (rRNA) genes (Ishihara et al., 2022).

In the present study, we report that Arabidopsis topoisomerase 1α gene (*TOP1α*) is cooperatively involved in leaf adaxialization together with *AS2-AS1*. Meta-analysis of microarray and genetic analyses revealed that repression of *KRP5/ICK3* gene, which encodes a member of the Cip/Kip family CDK inhibitor, is crucial for leaf adaxialization and formation of flat leaves. In addition, we found that the *top1α* mutation and treatment with 10-hydroxyl camptothecin (10H-CPT) changed the shape and pattern of AS2 bodies within the nucleolus, as seen in *rh10*, *nuc1,* and *rid2* mutant plants, in which nucleolar stress was induced (Ando et al. 2023). Furthermore, the phenotype of abaxialized filamentous leaves in the *as2 top1α* plants was suppressed by mutation of *SUPPRESSOR OF ROOT INITIATION DEFECTIVE TWO1* (*SRIW1*)/*ANAC082* which encodes a nucleolar stress response factor. We propose a novel role of AS2 as an attenuator to nucleolar stress, which acts to establish the adaxial-abaxial polarity of leaves and form flat leaves, even when the plant is exposed to nucleolar stress.

## RESULTS

### Camptothecin, a specific inhibitor of topoisomerase I, converts asymmetrically extended leaves of the *as2* mutant into a filamentous form

To elucidate the molecular network of leaf formation controlled by the *AS2* and *AS1* genes, 2,209 molecules containing natural compounds derived from Inter BioScreen Ltd, which are composed of microorganism and plant-derived natural compounds and their derivatives, were screened. In the first screening, Col-0 (wild type), *as2-1*, and *as1-1* plants were grown for 3 weeks at a concentration of 12.5 μM, and 35 molecules that showed growth inhibition were selected. In the second screening, 35 molecules were tested at various concentrations and 8 molecules were identified that produced filamentous leaves in *as2-1* and *as1-1* (Fig. 1A; Table 1).

**Figure 1.**
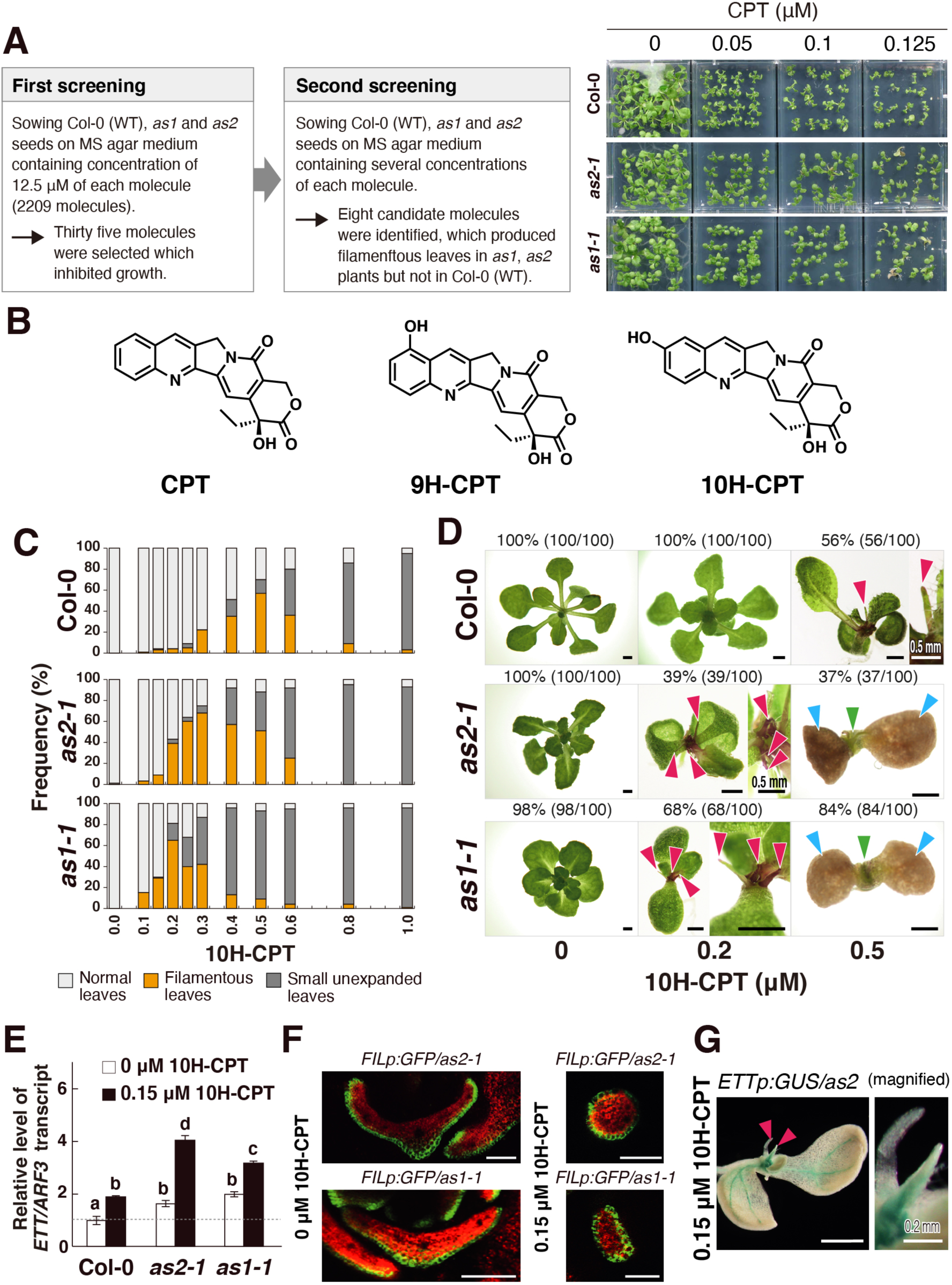
Procedures of screenings of molecules produced filamentous leaves in *as2-1* and *as1-1* mutants and characterization of phenotypes of plants treated with 10-hydroxycamptothecin (10H-CPT). (A) Left panel, experimental procedures for two-step screenings of molecules induced filamentous leaves in *as2-1* and *as1-1* mutants. Right panel, dose-dependent effects of CPT on leaf abnormalities and leaf formation. (B) Chemical structures of CPT, 9H-CPT, and 10H-CPT. (C) Dose-dependent effects of 10H-CPT on leaf abnormalities and leaf formation. Each value is calculated from the means of 3 biological replicates (n=100). (D) Phenotypes of Col-0, *as2-1*, and *as1-1* plants in the presence of DMSO, 0.2 and 0.5 μM 10H-CPT grown for 21days. Scale bars indicate 1 mm. Red arrowheads indicate filamentous leaves. Green arrowheads indicate small unexpanded leaves like clumps. Blue arrowheads indicate withered cotyledons. (E) Changes (mean ± SD) in transcription levels of the *ETT*/*ARF3* gene. Each value was normalized by reference to the level of *ACT2* transcripts. Values indicated by dashed lines are shown relative to the values for Col-0 plants (*p*<0.05, Tukey’s test after significant ANOVA). (F) *FILp:GFP*/*as2-1* and *as1-1* plants treated with DMSO and 0.15 μM 10H-CPT for 14 days. Scale bars = 100 μm. (G) *ETTp:GUS*/*as2-1* plant treated with 0.15 μM 10H-CPT for 14 days. Scale bars = 1 mm. Red arrowheads indicate filamentous leaves.

**Table 1.**
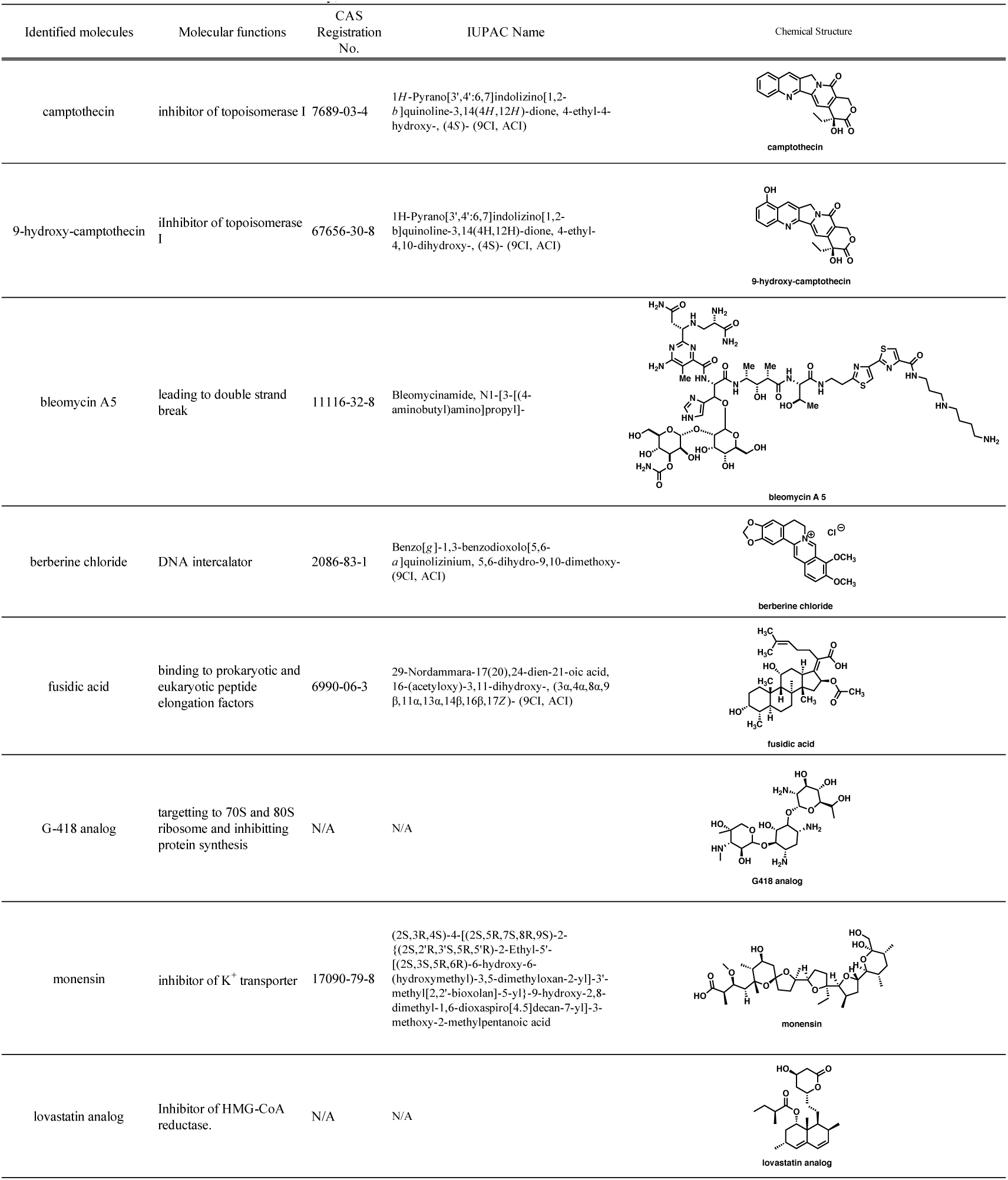
Molecules identified in the chemical library, which induced filamentous leaves in *as2-1* and *as1-1* mutants.

Identified 8 molecules and their related molecules were shown in Table 1 and Table 2. Four of the molecules (camptothecin, 9-hydroxyl camptothecin, bleomycin A5, and berberine chloride) are known as tumor suppressors in human cells, and inducers of nucleolar stress. Two of the molecules (fusidic acid and G-418 analog) are known as inhibitors in protein synthesis. Monensin is a molecule that has been reported to cause oxidative stress in animal cells. Lovastatin analogs, the primary target of which is the inhibition of HMG-CoA reductase (HMGR), the rate-limiting enzyme in cholesterol biosynthesis, are drugs used to treat dyslipidemia, and are molecules that suppress the synthesis of ATP, an essential energy source for cell survival. Statins are also known to be inhibitors of various tumors.

**Table 2.** Molecules identified in the chemical library, and its derivatives, which induced filamentous leaves in *as2-1* and *as1-1* mutants. The bold text indicates the chemical reagents selected by screening from the chemical library.

| Molecular characteristics | Substance | Comments | References |
| --- | --- | --- | --- |
| inhibitors of DNA topoisomerases | **camptothecin** | TOP I inhibitor, forming topoisomerase I-DNA covalent complex | Christensen et al., 2002; Rubbi et al., 2003; Basili et al., 2009; Ohwada et al., 2009; Burger et al., 2010; Boulon et al., 2010; Hutten et al., 2011; Grierson et al., 2013; Gonzalez-Arzola et al., 2024 |
|  | **9-hydroxy-camptothecin** |  |  |
|  | 10-hydroxy-camptothecin |  |  |
|  | etoposide | TOP II inhibitor, forming topoisomerase II-DNA covalent complex |  |
| DNA damage agents | **bleomycin A5** | leading to double strand break | Hecht, 2000; Palwai & Eriksson, 2011 |
|  | bleomycin A2+B2 |  |  |
|  | mitomycin C | inhibitor of DNA synthesis by introduction of cross-links into DNA at purine bases | Chan et al., 1988 |
|  | hydroxyurea | DNA replication inhibitor | Luong et al., 2018 |
|  | **berberine chloride** | DNA intercalators | Nakagawa et al., 2012; Avila-Carrasco et al., 2019 |
|  | coptisine |  |  |
|  | N-methylmesoporphyrin IX (NMM) |  |  |
|  | tricostatin A | epigenetic regulation, inhibitor of HDAC activity | Ueno et al., 2007 |
|  | 5-aza-2'-deoxycytidine | epigenetic regulation, inhibitors of DNA methylation | Iwasaki et al., 2013 |
| molecules targeting ribosomes | **fusidic acid** | binding to prokaryotic and eukaryotic peptide elongation factors | Kim et al., 2023 |
|  | **G-418** | targets to 70S and 80S ribosome and inhibits protein synthesis | Bidou et al., 2017; Heldin et al., 2023 |
|  | **G-418 analog** | N/A | N/A |
| inhibitor of K<sup>+</sup> transporter | **monensin** | inhibitor of K<sup>+</sup> transporter: inhibitor of Androgen signaling leading to Apoptosis in cancer cells | Ketola et al., 2010; Kocaturk et al., 2019 |
| inhibitor of HMG-CoA reductase | **lovastatin analog** | molecules that inhibits HMG-CoA reductase, a key regulator of metabolism, and are used as drug to reduce blood cholesterol and fat. | Nayan et al., 2017; Beckwitt et al., 2018; Lu, et al., 2018 |
|  | mevastatin |  |  |

Among them, in this study, we focused on CPT and 9-hydroxyl-CPT (9H-CPT), which inhibit specifically the action of type IB DNA topoisomerase (Fig. 1B). CPT was, however, too strong to control effects on leaf development. Since 9H-CPT was not commercially available, we used 10-hydroxyl-CPT (10H-CPT), which is less toxic than CPT (López-Meyer and Nessler, 1997; Lorence et al., 2004), and sought appropriate concentrations of 10H-CPT that affected leaf morphogenesis but not plant survival (Fig. 1C,D).

When *as2-1* and *as1-1* plants were grown on media supplemented with 10H-CPT (0.1-1.0 µM), most of the plants formed leaves with abnormal shapes, which were roughly classified into two groups (Fig. 1C,D): filamentous leaves (indicated by red arrowheads) and small unexpanded leaves like clumps (indicated by green arrowheads) between two browned cotyledons (indicated by blue arrowheads) in Fig. 1D. Even at low concentrations of 10H-CPT (0.2 µM), a half of these mutants generated filamentous or small unexpanded leaves, although almost all (95%<) the Col-0 plants generated normally expanded leaves under the same growth conditions (Fig. 1C,D). We decided to monitor leaf morphology of plants that had been grown in the presence of 0.15 and/or 0.2 µM 10H-CPT. We should also point that more than half of Col-0 plants produced filamentous leaves at 0.5 µM 10H-CPT (Fig. 1C,D).

We examined levels of transcripts of abaxial domain-determining gene *ETT/ARF3,* which is a direct downstream target of the AS2-AS1 complex (Iwasaki et al., 2013; Husbands et al., 2015), in shoot apices of 0.15 µM 10H-CPT-treated Col-0, *as2-1,* and *as1-1* plants. Real-time quantitative RT-PCR analysis revealed that levels of *ETT/ARF3* transcripts were higher in *as2-1* and *as1-1* than that in the Col-0 plants when treated with 10H-CPT (Fig. 1E). This result implies that the higher level of accumulation of transcripts of the abaxial factor *ETT/ARF3* by the treatment with 10H-CPT might be at least in part a reason for the generation of filamentous leaves.

To examine the adaxial-abaxial patterning in leaves of 10H-CPT-treated Col-0, *as2-1*, and *as1-1* plants, we used the *FILp:GFP* reporter gene in which the coding sequence for green fluorescent protein (GFP) was fused to the *FILAMENTOUS FLOWER* (*FIL*) promoter active in abaxial cells of wild-type leaves. We made transgenic plants (Col-0, *as2-1,* and *as1-1*) expressing the reporter gene and treated these transgenic plants with 10H-CPT to induce filamentous leaves. As shown in Fig. 1F, strong signals of the GFP fluorescence were detected in cells located at peripheral positions of the filamentous leaves induced by the treatment of *as2-1* and *as1-1* plants with 10H-CPT (upper and lower right panels), whereas signals were detected in cells located in the adaxial epidermis of the leaves of *as2-1* and *as1-1* plants without 10H-CPT (upper and lower left panels). Thus, the 10H-CPT treatment of *as2* and *as1* plants failed to develop the adaxial domain during leaf morphogenesis.

The GUS activity was observed in filamentous leaves of 10H-CPT-treated *as2* plants expressing *ETTp:GUS* (Fig. 1G). We analyzed GUS activity in filamentous leaves of 10H-CPT-treated *BREVIPEDICELLUS* (*BP*)*p:GUS/as2* plants, but GUS activity was not observed (Fig. S1B). The *bp-1 knat2-3 knat6-2* mutation did not suppress the filamentous leaf phenotype in 10H-CPT-treated *as2-1* mutants (Fig. S1A). These results suggested that the filamentous leaves caused in *as2-1* plants treated with 10H-CPT are due to increased *ETT/ARF3* expression, but not *BP*.

### *AS2* and *TOP1α* synergistically repress expression of abaxial determinant *ETT/ARF3* gene

An endogenous target molecule of CPT is TOPI, which loosens the super-coils and loop structures in double-stranded DNAs by mediating single-strand breakages and reunions. In the *A. thaliana* genome, TOPI proteins are encoded by two genes, *TOP1α* and *TOP1β,* which are tandemly arrayed (Fig. S2A) (Takahashi et al., 2002). We investigated genetic interactions among *AS2*, *AS1*, *TOP1α*, and *TOP1β*. The *as2-1 top1α-1* and *as1-1 top1α-1* double mutants exhibited phenotypes of the severely filamentous leaves (Fig. 2A, 2^nd^ row and Fig. S2B). Similar phenotypes were also observed in *as2-1 mgo1-7* and *as1-1 mgo1-7* double mutants (*mgo1-7* is another allele of *top1α*) (Figs. S2A,B,C; Graf et al., 2010). Mutants of *as2-1 top1β-1* and *as1-1 top1β-1* (or *top1β-2*), however, did not generate filamentous leaves (Fig. S2D). Thus, filamentous leaves are a synthetic phenotype specifically generated by double mutants, *as2-1 top1α-1* and *as1-1 top1α-1,* but not by *top1β*.

**Figure 2.**
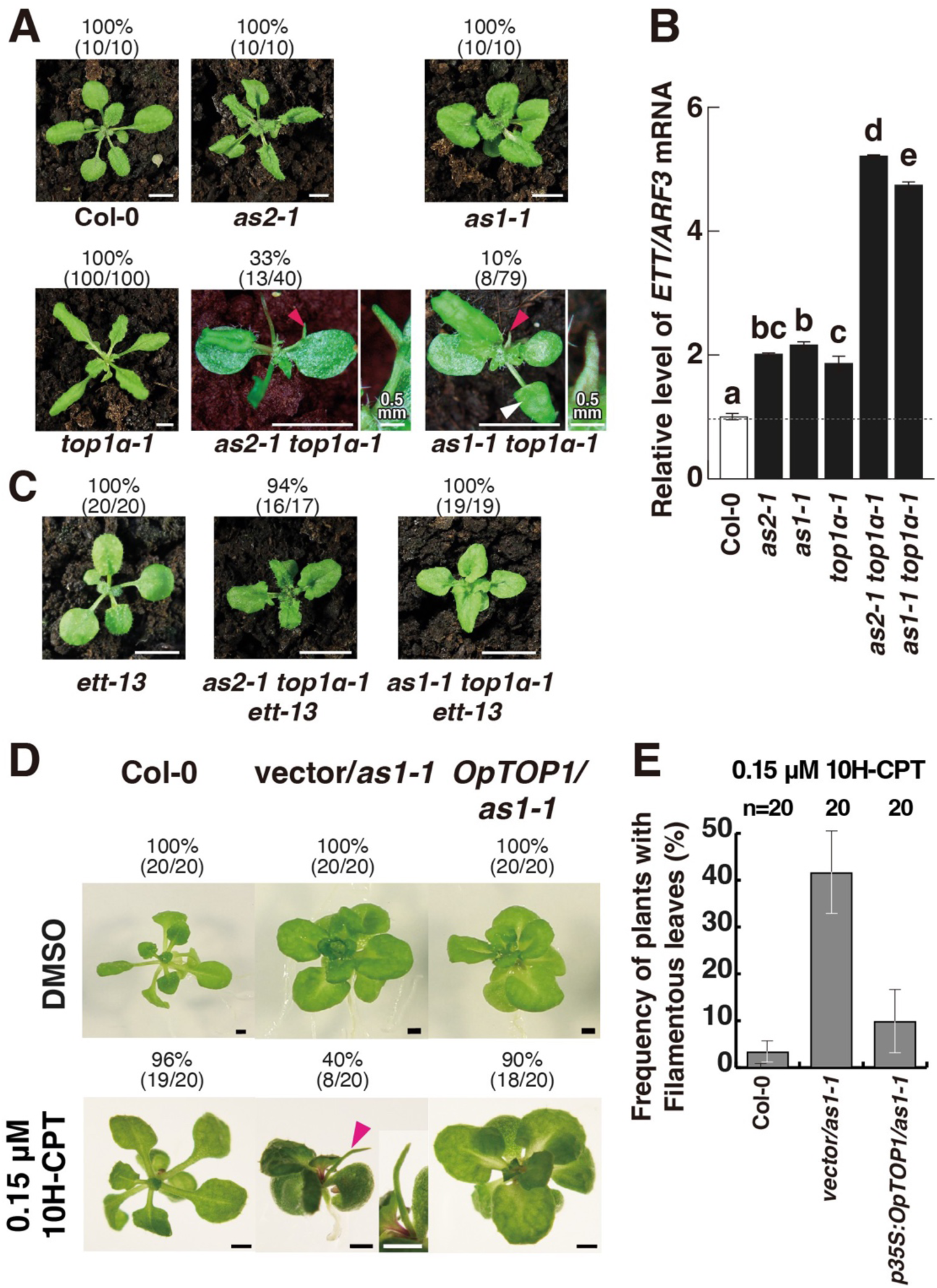
*TOP1α* and *AS2-AS1* cooperatively function in development of expanded leaves with the proper adaxial-abaxial polarity. (A) The mutation of *TOP1α* exhibited impairment of leaf adaxial/abaxial polarity in *as2-1* and *as1-1* backgrounds. Leaf phenotype of the plants at 21 or 22 days after sowing (DAS). Bars = 5 mm (unless otherwise indicated). Red and white arrowheads indicate filamentous and trumpet-like leaves, respectively. (B) Changes (mean ± SD) in transcription levels of the *ETT*/*ARF3* gene. Each value was normalized by reference to the level of *ACT2* transcripts. Values indicated by dashed lines are shown relative to the values for Col-0 plants (*p*<0.05, Tukey’s test after significant ANOVA). (C) Suppression of leaf phenotypes of *as2-1 top1α-1* by *ett-13*. Leaf phenotype of the mutant plants at 22 DAS. Bars = 10 mm. (D) The *OpTOP1* gene derived from CPT-producing herbs, *Ophiorrhiza pumila,* attenuates the action of 10H-CPT to induce the leaf filaments in Arabidopsis. Comparison of leaf morphology among Col-0, empty vector-introduced *as1-1*, and *as1-1* expressing *OpTOP1* plants growing in the absence or the presence of 10H-CPT. (E) Quantitative analyses of frequencies of filamentous leaves in plants of Col-0, *as1-1* transformed with empty vector pGWB2, and *as1-1* expressing *OpTOP1*. Data represent mean ±SD of 3 replicates (n=20 for each).

Since the *ETT/ARF3* gene is a direct target of the AS2-AS1 complex (Iwasaki et al., 2013; Husbands et al., 2015), we examined levels of transcripts in *top1α-1, as2-1 top1α-1*, and *as1-1 top1α-1*. As shown in Fig. 2B, RT-PCR analysis revealed that levels of *ETT/ARF3* transcript were higher in the shoot apices of these double mutants than those in Col-0 plants and each single mutant (*top1α-1, as2-1,* and *as1-1*). The above results suggest that *TOP1α* and *AS2-AS1* synergistically repress transcript levels of the *ETT/ARF3* gene, the abaxial-determinant, which leads to the development of the adaxial domain of leaves. We examined whether increased levels of *ETT/ARF3* transcripts might be responsible for the formation of filamentous leaves in *as2-1 top1α-1* and *as1-1 top1α-1* (Fig. 2A) by introducing the *ett-13* mutation into each of the double mutants. As shown in Fig. 2C, the *ett-13* mutation suppressed the filamentous leaf phenotype in these double mutants. These results suggest that the increase of levels of the *ETT/ARF3* expression in *as2-1 top1α-1* and *as1-1 top1α-1* plants causes the formation of filamentous leaves. In Col-0, repression of *ETT/ARF3* by a cooperative action of *TOP1α* and *AS2-AS1* is critical in the leaf development with adaxial-abaxial polarity.

These results, together with those of the previous section with 10H-CPT, imply that a synergistic action of *AS2* (and *AS1*) and *TOP1α* might play an essential role in development of flat and expanded leaves in Arabidopsis.

### *Op*TOP1 from the camptothecin-producing herb confers camptothecin resistance to Arabidopsis plants

CPT was originally discovered in a medical plant, *Camptotheca acuminata,* and is known to be produced by many species of *Ophiorrhiza* including *Ophiorrhiza pumila* (Sirikantaramas et al., 2008). TOPI proteins of *Ophiorrhiza* plants (*Op*TOP1) have mutated amino acid residues in the region that are responsible for the catalytic function or CPT binding as compared to those of *A. thaliana* and *Homo sapiens*. Yeast cells expressing the *OpTOP1* cDNA from *Ophiorrhiza pumila* confer higher resistance to CPT than those expressing *TOP1* cDNAs of *A. thaliana* and *H. sapiens.* The 35S promoter-fused *OpTOP1* cDNA was constructed, introduced into the *as1-1* mutant, and tested for effects of 0.15 μM 10H-CPT on the formation of filamentous leaves in these transformed plants. Fig. 2D showed that 90% of the *as1-1* mutants expressing *OpTOP1* formed normally expanding leaves in the presence of 0.15 μM 10H-CPT, although approximately 40% of *as1-1* transformed with empty vector pGWB2 formed filamentous leaves at the same concentration of 10H-CPT (Fig. 2D,E). These results suggest that *OpTOP1* in the *as1-1* mutant is functionally replaced with activity of endogenous TOP1 of Arabidopsis that was sensitive to 0.15 μM 10H-CPT. Thus, Type I topoisomerase plays a critical role in the formation of a flat and symmetrically expanding leaves.

### *AS2* and *TOP1α* act synergistically to repress the transcript levels of the CDK inhibitor *KRP5/ICK3*, which is important for the formation of normally expanded leaves

We previously proposed that transcription of *KRP2/ICK2* and *KRP5/ICK3* genes, which encode CDK inhibitors involved in the progression of cell division cycle, is regulated downstream of AS2-AS1 (Takahashi et al., 2013). In the present experiment, we analyzed patterns of gene expression data from DNA microarrays using mRNAs from shoot apices of the 10H-CPT-treated *as2-1* and Col-0 plants, and extracted genes regulated downstream of AS2, which were classified into 30 clusters by using the knowledge-based fuzzy adaptive resonance theory (KB-FuzzyART) (Takahashi et al., 2003; 2008; 2013) (Fig. S3A; Table S1).

Expression levels of genes which were classified to clusters 10, 12, 16, and 25 (Figs. 3A, S3A,B,C) were increased in both *as2-1* and Col-0 treated with 10H-CPT. Genes involved in ‘cell fate’ (such as *BP* and *ETT*/*ARF3*) were significantly enriched in the clusters 10, 12, 16, and 25 (Figs. S3A, S3B; Tables S2-S4). Therefore, we focused on those 556 genes in clusters 10, 12, 16, and 25 (Fig. 3A).

**Figure 3.**
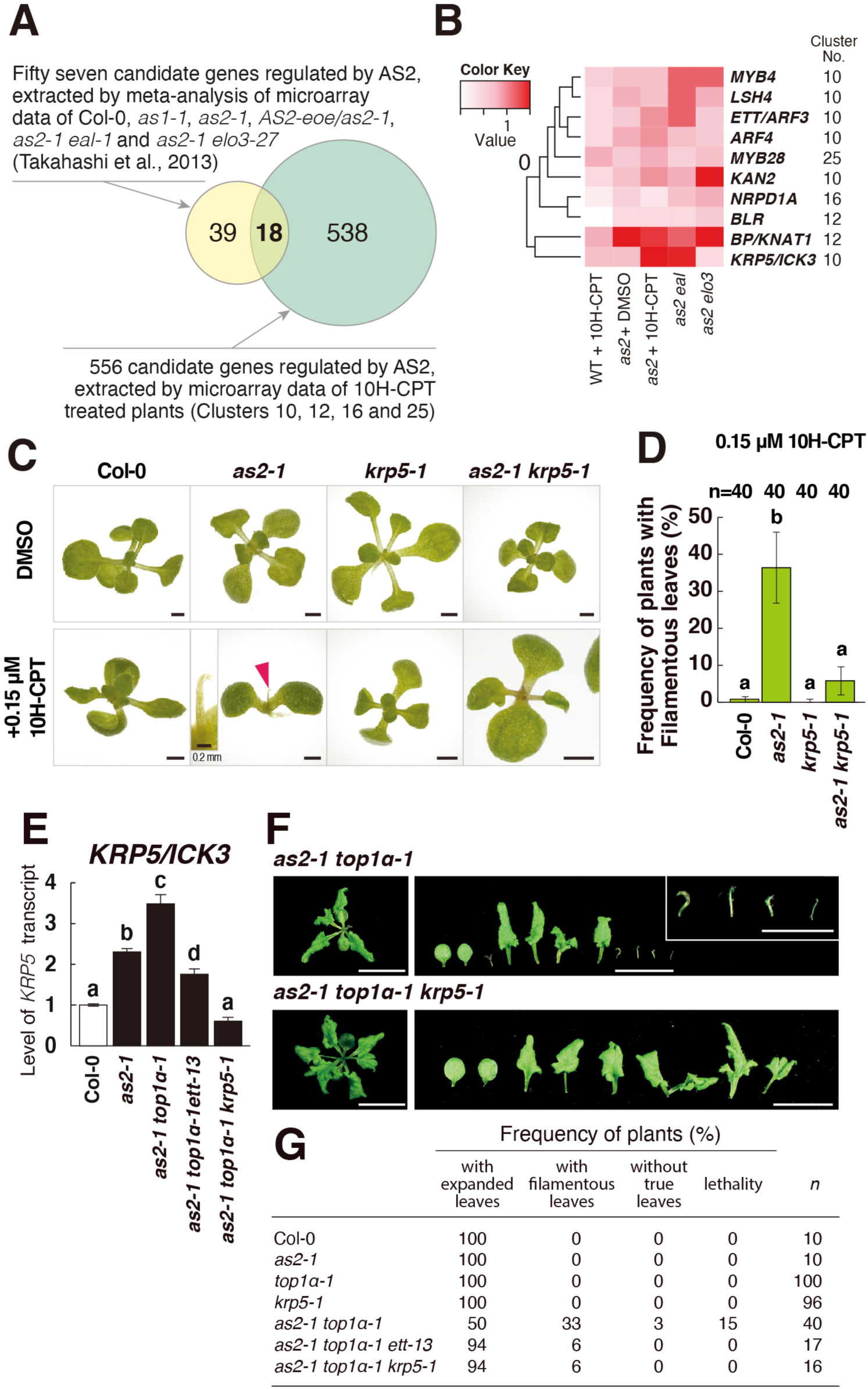
The *KRP5/ICK3* gene is the downstream gene regulated by *AS2-AS1* and *TOP1α.* (A) Venn diagram that shows the relationship between 57 candidates regulated by *AS2*, *AS1*, *ELO3* and *EAL* (Takahashi et al., 2013) and 556 genes in clusters 10, 12, 16, and 25 extracted in this experiment (Figure S3). Eighteen genes are overlapped. (B) Heatmap representing transcript levels of 10 genes that are candidates for being downstream of AS2-AS1. (C) The *krp5-1* mutation suppressed filamentous leaf phenotype in *as2-1* treated with 0.15 μM 10H-CPT for 14 days. Bars = 1 mm. A red arrowhead indicates a filamentous leaf. (D) Frequencies of plants with filamentous leaves at 1^st^ and 2^nd^ leaf positions when plants were treated with 0.15 μM 10H-CPT Data represent mean ±SD of 3 replicates (n=40 for each). (*p*<0.05, Tukey’s test after significant ANOVA). (E) Changes in levels of transcript of *KRP5/ICK3* in shoot apices. Each value was normalized by reference to the level of *EF1α* transcripts (*p*<0.05, Tukey’s test after significant ANOVA). (F) Gross morphology of *as2-1 top1α-1*, *as2-1 top1α-1 krp5-1.* Bars =10 mm (except the bars in inset of *as2-1 top1α-1* are 5 mm). Plants photographed at 22 DAS. (G) Frequencies of phenotypes of leaves and plant lethality at 22 DAS.

Next, to extract common downstream candidate genes involved in formation of filamentous leaves we identified genes common to 57 candidate genes regulated by both *BOBBER1* (*BOB1*)/*ENHANCER of AS2 and AS1* (*EAL)* and *ELONGATA3* (*ELO3*)/*ENHANCER of ASYMMETRIC LEAVES TWO1*(*EAST1*), which have been shown to act as modifiers of *as2-1* and *as1-1* (Fig. 3A; Kojima et al., 2011; Ishibashi et al., 2012; Takahashi et al., 2013).

The group of 18 selected genes included 10 genes encoding proteins with known domains such as *ETT*/*ARF3* and *BP*, among which *KRP5/ICK3*, a Cip/Kip family CDK inhibitor gene, was included as a gene in the cluster of ‘division-related genes’ (Fig. 3B, S3D; Table S1). Quantitative RT-PCR analysis confirmed that the level of *KRP5/ICK3* transcripts was obviously increased in the shoot apex of 10H-CPT-treated *as2-1* plants (Fig. 3E). The transcription levels of the other six *KRP* genes did not increase in *as2-1* treated with 10H-CPT, indicating that *KRP5/ICK3* is the only common downstream gene among the *KRP* genes (Figs. S3E, S3F).

Treatment of the *as2-1* plant with 10H-CPT gave rise to the production of filamentous leaves in 35% of the plants (Fig. 3C,D). The efficiency of occurrence of such filamentous leaves in the *as2-1 krp5-1* mutant was reduced to only 5% (Fig. 3C,D). Treatment of *as2-1 krp5-4* with 10H-CPT similarly reduced the efficiency of formation of filamentous leaves (Fig. S4). While, treatment of *as2-1 krp2-5* with 10H-CPT did not reduce the efficiency of formation of filamentous leaves (Fig. S5). Thus, negative regulation of *KRP5/ICK3* should be necessary for the formation of normally expanded leaves.

Next, we measured levels of *KRP5/ICK3* transcripts in *as2-1* single and *as2-1 top1α-1* double mutants. As shown in Fig. 3E, levels of *KRP5/ICK3* transcripts in the double mutant were higher than *as2-1* single mutants and introduction of *ett-13* into the *top1α-1 as2-1* double mutant decreased the level of the *KRP5/ICK3* transcript, suggesting that repression of *KRP5/ICK3* is controlled synergistically by *AS2* and *TOP1α* through the *ETT*/*ARF3* function.

We examined whether filamentous leaf phenotypes of the *as2-1 top1α-1* double mutant might be suppressed by introduction of the *krp5-1* mutation. As shown in Fig. 3F and 3G, the introduction of the *krp5-1* mutation into the double mutant decreased the proportion of plants showing the filamentous leaves (from 33% to 6%).

Since *KRP5/ICK3* is a repressive regulator of the cell division cycle, *AS2* and *TOP1α* might be positively involved through repression of expression of *KRP5/ICK3* in the development of the adaxial domain at least in part through some steps of progression of cell division cycle.

### *AS2* and *TOP1*α synergistically control proliferation of cells around the shoot apical meristem

We measured relative levels of *KRP5/ICK3* transcripts during cell cycle progression of synchronized cultured cell line MM2d of *A. thaliana* (Fig. 4A). The transcript levels of *KRP5/ICK3* and *HISTONE H4* (a S phase marker) peaked at 2-4 hours after the removal of aphidicolin (corresponding to S phase), being consistent with previous microarray results (Menges et al., 2002). The broad pattern of transcripts of *KRP5/ICK3* was also found during the progression of M phase, overlapping with peaks of the transcripts of M-phase-specific genes such as kinesin *AtNACK1/HINKEL* (Nishihama et al., 2002; Strompen et al., 2002) and *KNOLLE* (Heese et al., 2001). These findings were consistent with the notion that *KRP5/ICK3* might act as a factor in a repressive manner on the progressions of S and M phases of the cell cycle.

**Figure 4.**
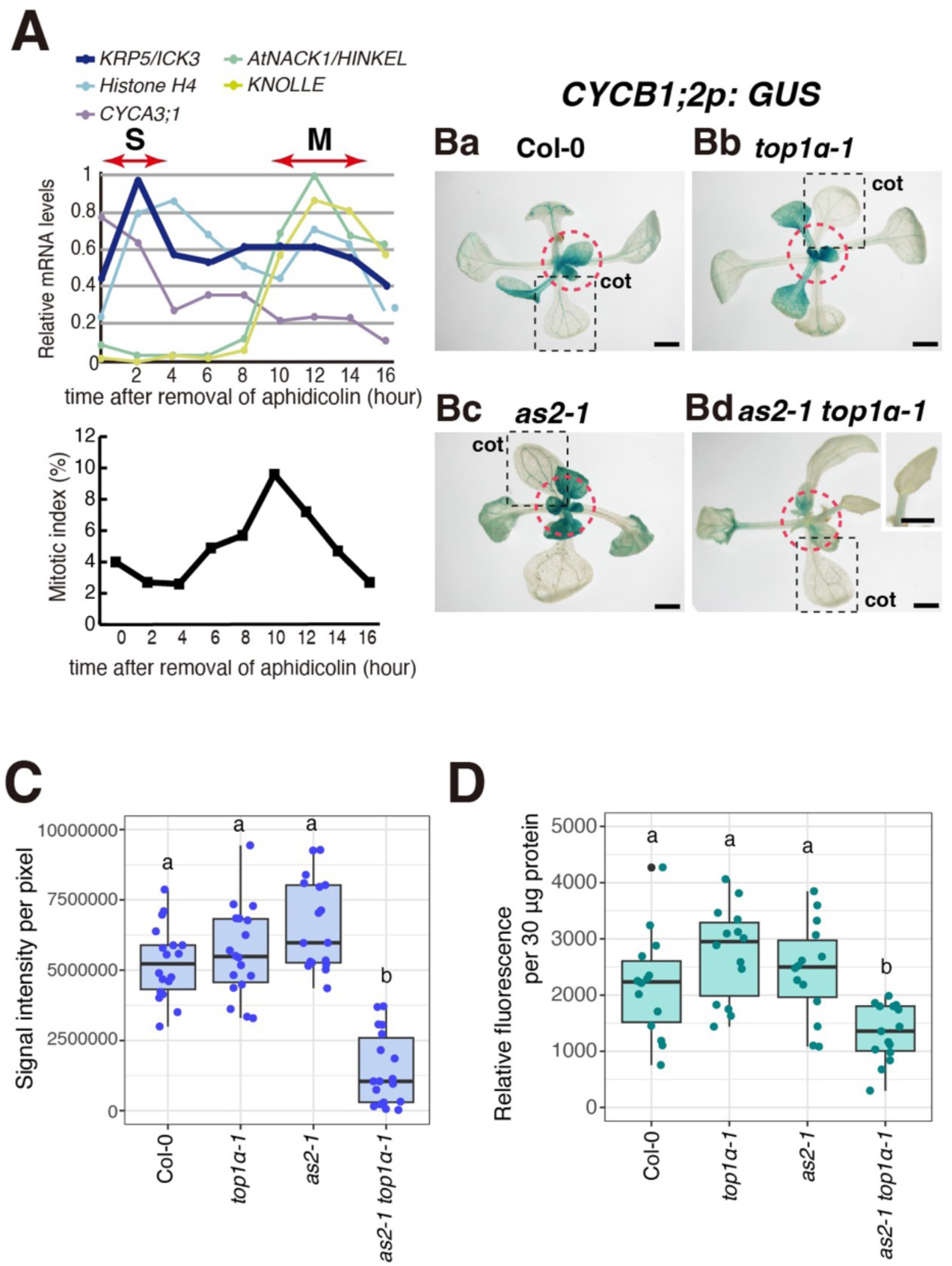
Relative levels of *KRP5/ICK3* transcripts during cell cycle progression and cell division potential of leaves in *as2-1 top1α-1.* (A) Transcript levels of the *KRP5/ICK3* gene (upper panel) and mitotic index (lower panel) in aphidicolin-synchronized Arabidopsis MM2d cells. (B) Representative GUS expression patterns of 13-day-old Col-0 (Ba), *top1α-1* (Bb), *as2-1* (Bc) and *as2-1 top1α-1* (Bd) mutant plants that harbored the *CYCB1;2p:GUS*. Shoot apices are marked by dashed red circles and cotyledons are marked by dashed black squares. Bars = 1 mm. (C) Relative GUS activity around shoot apices using X-Gluc calculated by ImageJ. One-way ANOVA followed by Tukey’s HSD analysis was performed. Letters (a, b) indicate that the signal intensities are statistically different within the group. (*P*<0.01) n = 18. (D) Relative GUS activity from aerial part using 4-MU. One-way ANOVA followed by Tukey’s HSD analysis was performed. Letters (a, b) indicate that the fluorescence is statistically different within the group. (*P*<0.01) n = 15.

Using transgenic plants expressing the *CYCB1;2p:GUS* gene as a marker that is expressed at early M phase (Ishibashi et al., 2012), we examined the cell division activity of local regions in shoot apices that cover developing leaves of wild type (Col-0), *top1α-1*, *as2-1*, and *as2-1 top1α* plants (marked by dashed red circles). As shown in Figs 4Ba, Bb, and Bc, clear GUS activities were detected as blue staining in these regions. However, only a limited level of GUS activity was detected in shoot apices of the double mutant (marked by the dashed red circle in Fig. 4Bd). Fig. 4C and 4D showed relative GUS activity in shoot apices that quantified by analyzing X-gluc staining using ImageJ and that in aerial part using 4-MU, respectively. Levels of GUS activities in *top1α-1 as2-1* were clearly decreased (Fig. 4C,D). These data suggest that the cell division activity of cells of the shoot apices in *as2-1 top1α-1* is limited as compared to those in the wild type plant, corresponding to the increase in the level of transcript of the *KRP5/ICK3* gene in *as2-1 top1α-1* (Fig. 3E).

Therefore, we conclude that *AS2* and *TOP1α* genes synergistically and positively control proliferation of cells of shoot apices by repression of *KRP5/ICK3*, which is critical for formation of flat and expanded leaves.

### Chemical inhibitors of topoisomerase II and DNA replication induce formation of filamentous leaves in the *as2* mutant background

As shown in Fig. 5A and Fig. S6A, we here further tested following reagents for effects on the leaf development on *as2-1* and *as1-1* backgrounds: etoposide, a specific inhibitor of type II DNA topoisomerase (TOPII), which induces a double-stranded break, allows the passage of a double-stranded DNA segment and rejoins the cleaved DNAs; mitomycin C, an inhibitor of DNA synthesis by introduction of cross-links into DNA at purine bases; bleomycin A5, which leads to double strand break. The *as2-1* and *as1-1* mutant plants that were grown in the presence of these molecules formed filamentous leaves, although Col-0 plants did not, as observed upon the treatment with 10H-CPT (Fig. 5A; Fig. S6A).

**Figure 5.**
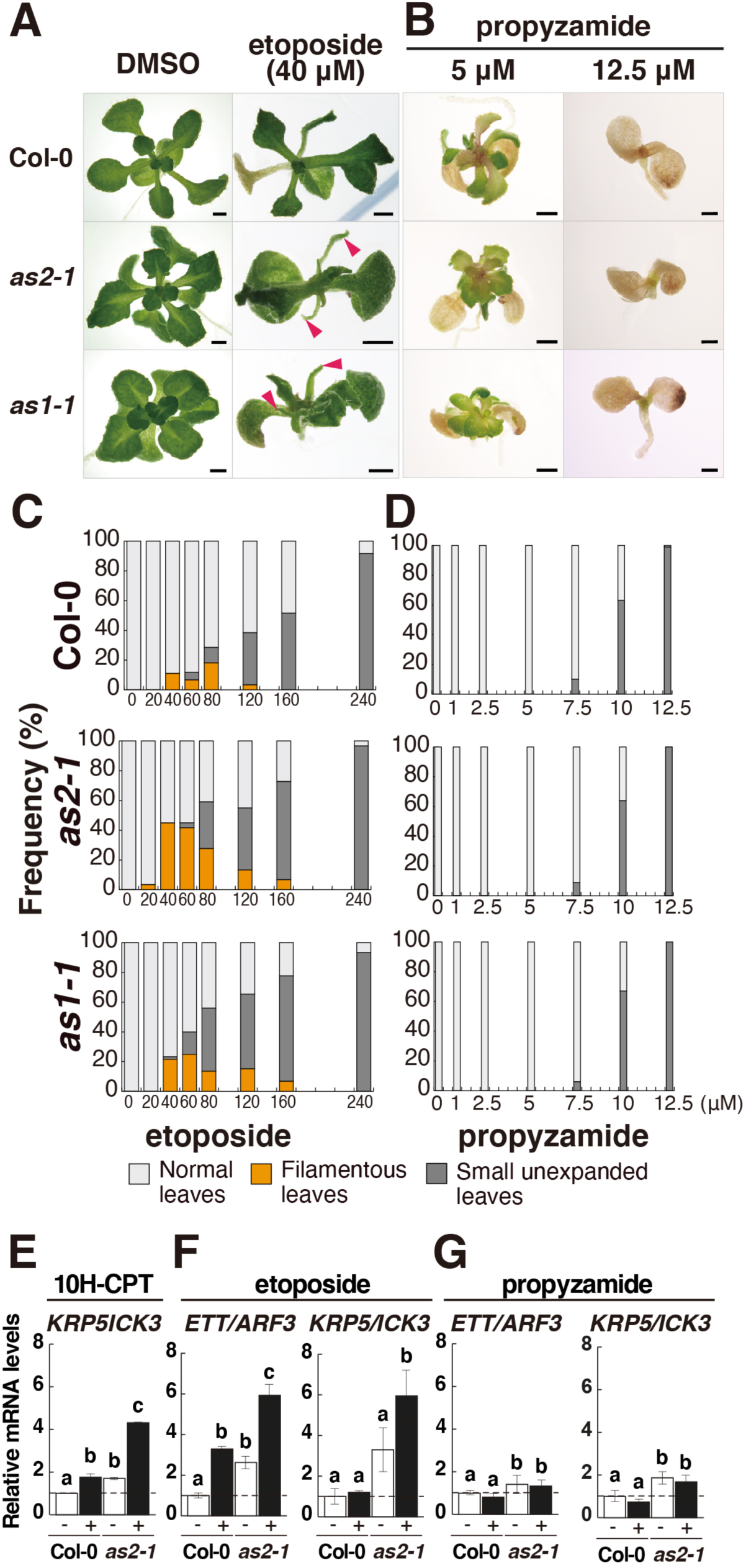
Inhibitors of topoisomerase II induce formation of filamentous leaves, but not those with inhibitors of microtubule turnover. (A) Phenotypes of Col-0, *as2-1*, and *as1-1* plants in the presence of DMSO, etoposide grown for 21days. Red arrowheads indicate filamentous leaves. Bars = 1 mm. (B) Representative phenotypes of plants treated with 5 and 12.5 μM propyzamide for 21 days. Bars = 1 mm. (C, D) Dose-dependent effects of etoposide (C) and propyzamide (D) on leaf abnormalities and leaf formation. Each value is calculated from the means of 3 biological replicates (n=20 for etoposide and n=24 for propyzamide), respectively. (E) Changes (mean ± SD) in transcription levels of *KRP5/ICK3* in shoot apices Col-0 and *as2-1* treated without and with 10H-CPT. (F, G) Changes in transcription levels of *ETT*/*ARF3* and *KRP5/ICK3* in shoot apices of plants treated without and with 40 μM etoposide (F), and 5 μM propyzamide (G) (*p*<0.05, Tukey’s HSD test after significant ANOVA).

These results implicate that the normal progression of DNA replication during the cell division cycle and/or the maintenance of appropriate topological features of DNA molecules during canonical DNA replication, repair, and recombination are critical for the development of flat and expanding leaves.

The progression of the M phase of the cell cycle is supported by the metabolism of microtubules by their dynamic instability. We subsequently examined effects of reagents such as propyzamide, oryzalin, and paclitaxel that inhibit turnover of microtubules, which inhibits the progression of M phase of cell cycle at prometaphase (Kakimoto and Shibaoka, 1988; Nishihama et al., 2001; Nishihama et al., 2002; Sasabe et al., 2011). Upon the treatments with these inhibitors, only dose-dependent growth inhibition was observed, but no filamentous leaves were generated (Fig. 5B,D; Fig. S6).

We next measured levels of *ETT/ARF3* and *KRP5*/*ICK3* transcripts in *as2-1* plants that had been treated with etoposide and propyzamide. The results showed that the levels of *ETT/ARF3* and *KRP5*/*ICK3* transcripts significantly increased in shoot apices of 10H-CPT, etoposide-treated *as2-1* plants (Fig. 5E,F). However, levels of transcripts of both genes did not further increase in the propyzamide-treated *as2-1* plant (Fig. 5G). Thus, there were tight relationships among the inhibition of TOPI and TOPII, the formation of filamentous leaves and the increases in *ETT*/*ARF3* and *KRP5*/*ICK3* transcripts.

### AS2 attenuates nucleolar stress induced by treatment with 10H-CPT and loss of function of *TOP1α* through the subnucleolar patterning of AS2 bodies

Cell biology with the recombinant AS2-fused YFP protein (yellow fluorescent protein: AS2-YFP) has shown that the AS2 protein mostly appears as two speckle-like structures at periphery of nucleolus, named AS2 bodies (Ueno et al., 2007; Luo et al., 2012; 2020; Ando et al., 2023).

We have reported that nucleolar proteins such as NUC1, RH10, and RID2 are involved in the subnucleolar patterning of AS2 bodies (Ando et al., 2023). It has also been reported that structural and functional defects in the nucleolus are observed in *nuc1*, *rh10*, and *rid2* mutants, leading to induction of nucleolar stress (Ohbayashi et al., 2017; Ohbayashi & Sugiyama 2018). We have examined whether 10H-CPT could affect the morphology and number of AS2 bodies in the nucleoli as described in Materials and Methods. As shown in Fig. 6A, the number of AS2 bodies markedly increased and their shapes of bodies were heterogeneity, as the concentration of 10H-CPT increased in the cultures.

**Figure 6.**
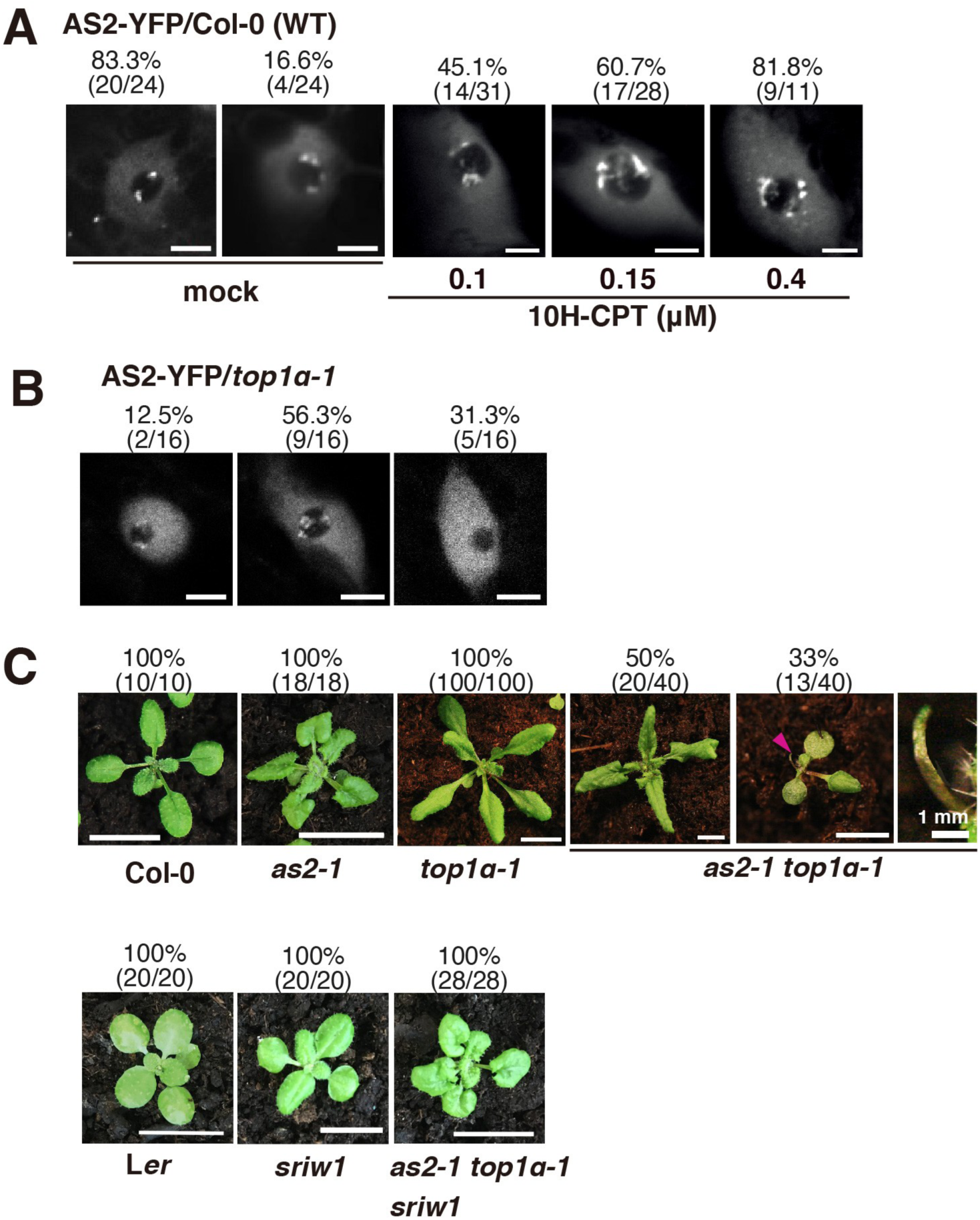
Treatment with 10H-CPT and *top1α* mutation affects subnucleolar patterning of AS2 bodies and the *sriw1* mutation suppresses leaf morphology in *as2-1 top1α-1.* (A) Images showing the signals from AS2-YFP fusion proteins in Col-0 (wild-type) plants, treated with DMSO (mock), 0.1, 0.15, or 0.4 μM 10H-CPT. (B) Images showing the signals from AS2-YFP fusion proteins in the *top1α*-*1* mutant. YFP-positive nuclei in (A) and (B) were observed in cells of the adaxial epidermis of cotyledons. Photographs in (A) and (B) show different types of subnucleolar patterning of AS2 bodies. Numbers above each photo show numbers of cells with AS2 body patterning exemplified by the photo/numbers of cells examined. Bars = 5 μm. (C) Suppression of leaf phenotypes of *as2-1 top1α*-*1* by *sriw1,* which has mutation in *ANAC082*. Leaf phenotype of the plants at 22 DAS. Bars =10 mm. Photographs show different types of leaf morphology. Numbers above each photo show numbers of plants with leaf morphology exemplified by the photo/numbers of plants examined.

We next examined effects of the *top1α-1* mutation on morphology of AS2 bodies. The number of AS2 bodies increased in 56% (9/16) cells of the *top1α-1* mutant, and obvious bodies were not observed in 31% (5/16) cells of the mutant (Fig. 6B). Therefore, TOP1α might be required for formation and localization of two AS2 bodies in the interior and periphery of the nucleolus. These results are similar to those observed in the *nuc1*, *rh10*, and *rid2* mutants (Luo et al., 2020; Ando et al., 2023).

It has been reported that the *sriw1* mutant, which has a mutation in the *ANAC082* gene involved in nucleolar stress response, suppresses the narrow leaves phenotype of the *rid2* and *rh10* mutant (Ohbayashi et al., 2017; Ohbayashi and Sugiyama 2018). The *top1α-1* mutant formed narrow leaves as observed in *rid2* and *rh10* plants (Fig. 2A; Fig. 6C). To determine which leaf phenotypes of *top1α-1 as2-1* were suppressed by *sriw1*, we next generated *top1α-1 as2-1 sriw1* triple mutant. The triple mutant did not exhibit a filamentous leaf phenotype, and displayed a phenotype nearly identical to that of *as2-1* (Fig. 6C). These results showed that the *sriw1* mutation suppressed *top1α-1 as2-1* mutation phenotype, such as filamentous leaves, but did not suppress *as2* mutation phenotype, suggesting that the AS2-dependent regulation of cell proliferation and development is not mediated by ANAC082. These results also indicated that nucleolar stress is induced in the *top1α* mutant. AS2 may function as an attenuator of deficiency in formation of leaves under nucleolar stress. The formation of AS2 bodies in nucleolus might be related to protection against nucleolar stress.

## DISCUSSION

### AS2 functions not only as a leaf morphogenetic regulator but also as a factor that alleviates nucleolar stress through the formation of AS2 bodies

In this study, we report that the *AS2* and *TOP1α* genes play a crucial role in cooperatively repressing the transcription of the abaxial-determinant *ETT/ARF3* gene. (Fig. 2). This regulation is essential for the development of left-right symmetry expanded leaves (Iwasaki et al., 2013; Husbands et al., 2015). In addition, dysfunction of *TOP1α* disrupts the shape and pattern of AS2 bodies within the nucleolus (Fig. 6; Fig. 7A). AS2 bodies partially overlap with chromocenters containing NORs, which consist of condensed and packed heterochromatin from 45S rDNA repeats (Fig. 7A; Luo et al., 2020). Formation of AS2 bodies is essential for AS2 function, as the YFP-fused *as2* mutant protein fails to form nucleolar bodies with wild-type morphology, resulting in abnormal leaf development (Luo et al., 2020; Ando et al., 2023).

**Figure 7.**
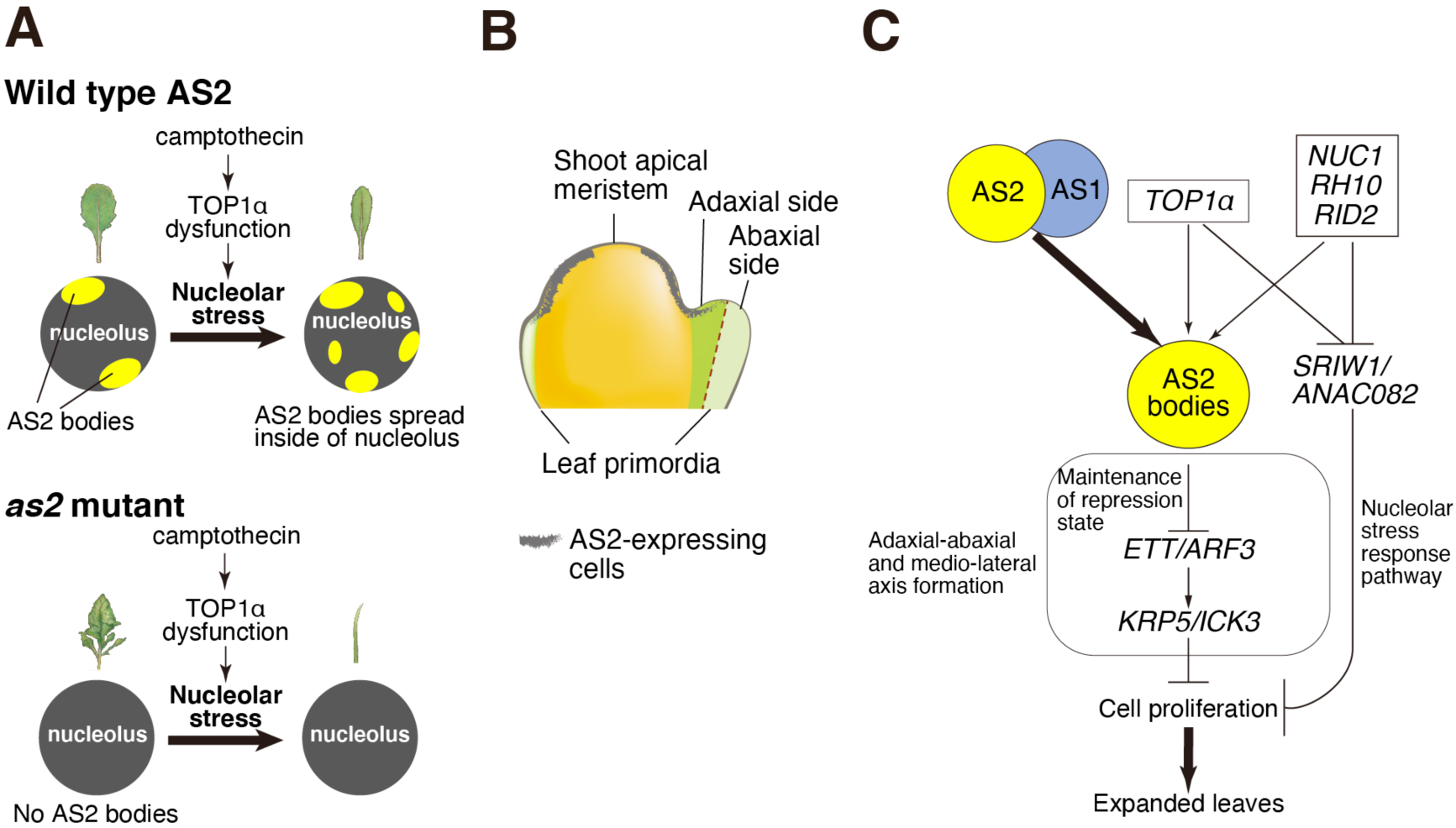
Roles of AS2, TOP1α and nucleolar proteins in shoot apices during the formation of expanded flat leaves. (A) Leaf phenotypes and localization of AS2 bodies in Col-0 and *as2*, when nucleolar stress was induced. (B) Developmental compartments in the shoot apex around the apical meristem are schematically shown. (C) Roles of the AS2–AS1 complex, TOP1α, and nucleolar proteins (NUC1, RH10, and RID2) in the regulation of *ETT*/*ARF3* and *KRP5/ICK3* genes via AS2 bodies in early stages of leaf primordia for the formation of expanded flat leaves. Nucleolar stress response via *SRIW1*/*ANAC082* is shown.

Based on these findings, we propose that AS2 and TOP1α proteins regulate rDNA repeat condensation through interactions with the NORs chromocenter via AS2 bodies. Additionally, these proteins might influence rDNA transcription by RNA polymerase I, ensuring appropriate ribosomal RNA levels. As noted in the Introduction, mutations in *RH10*, *RID2*, and *NUC1*, which encode the nucleolar proteins, all involved in nucleolar function, similarly affect AS2 body nucleolar patterning (Kojima et al., 2007; Pontvianne et al., 2007; Ohbayashi et al., 2011; Matsumura et al., 2016; Luo et al., 2020; Ando et al., 2023).

Previous *in vivo* and *in vitro* studies have demonstrated that TOPI is involved in rDNA transcription by RNA polymerase I in animal and yeast cells (Zhang et al., 1988). In animals, CPT and its derivatives, such as topotecan, inhibit TOPI and induce nucleolar stress, leading to defects in ribosome biogenesis (Mo et al., 2002; Lafita-Navarro & Conacci-Sorrell, 2023). Moreover, dysfunction of topoisomerases, including TOPI and TOPII, has been associated with nucleolar stress and p53-mediated responses in animals (Boulon et al., 2010; Ishihara et al., 2022). Meanwhile, the filamentous leaf phenotype caused by defects in *TOP1α* was suppressed by a mutation in *SRIW1*/*ANAC082*, a gene implicated in the nucleolar stress response of plant cells (Fig. 6C). Although the mechanisms of nucleolar stress response differ between animals and plants, they share the common feature that dysfunction of topoisomerases causes nucleolar stress.

Furthermore, our results demonstrate that the 10H-CPT-treated *as2-1* mutant and the *as2-1 top1α-1* double mutant exhibit significantly more severe phenotypes than either 10H-CPT-treated Col-0 or the *top1α* single mutant (Fig. 1D; Fig. 2A). This suggests that wild-type AS2 mitigates abnormalities induced by 10H-CPT treatment or *top1α* mutation. Thus, AS2 functions not only as a morphogenetic regulator but also as a factor that alleviates nucleolar stress, contributing to stable plant growth under fluctuating environmental conditions. The interplay between AS2-mediated gene repression and nucleolar stress mitigation remains to be further explored through molecular, genetic, and epigenetic analyses.

### The repression of *KRP5/ICK3* by *AS2* is a crucial process in the formation of flat leaves

We demonstrated that the *as2 top1α* double mutants developed abaxialized filamentous leaves, accompanied by elevated *KRP5/ICK3* expression (Fig. 3E). In addition, *krp5* mutations suppressed the formation of these filamentous leaves, while *CYCB1;2p:GUS* activity was reduced in the *as2 top1α* double mutants (Figs. 3; Fig. 4). Together with previous findings (Takahashi et al., 2013), these results suggest that AS2 represses *KRP5/ICK3* expression via *ETT*/*ARF3*, thereby ensuring proper cell cycle progression in adaxial cells during early leaf development (Fig. 7C).

Arabidopsis harbors seven *KRP* genes that belong to the Cip/Kip family of CDK inhibitors—comparable to the mammalian p21, p27, and p57 (De Veylder et al., 2001). CDKs promote cell cycle progression by inhibiting the RETINOBLASTOMA-RELATED 1 (RBR1) protein, which itself impedes G1 phase progression (Desvoyes & Gutierrez, 2020). In the root meristem, KRP5/ICK3 helps maintain meristem and stem cell activity and is a target of PLETHORA2, a factor known to inhibit G1 progression (Santuari et al., 2016). Recent studies indicate that the G1 phase is shortened in *krp5* mutants, implicating both KRP5/ICK3 and RBR1 as critical negative regulators of G1 duration in the root apical meristem (Echevarria et al., 2022). These regulatory mechanisms may also operate in the shoot apex, suggesting that KRP5/ICK3 could similarly suppress cell cycle progression in shoot apex cells. Consequently, the repression of *KRP5/ICK3* gene expression by *AS2* and *AS1* appears essential for promoting cell cycle progression following adaxial cell differentiation during early leaf formation (Fig. 7B,C).

The AS2-AS1 complex likely facilitates rapid cell division in early leaf primordia by suppressing *KRP5/ICK3* expression through the *ETT*/*ARF3* pathway (Fig. 7B,C). It is conceivable that the *ETT*/*ARF3* gene locus is recruited to AS2 bodies at the nucleolar periphery in adaxial cells during early leaf development, leading to its stable repression (Fig. 7B,C). Future studies should focus on elucidating the epigenetic mechanisms underlying *ETT*/*ARF3* suppression.

### AS2 is a guardian angel for alleviating nucleolar stress and promoting flat leaf development

The nucleolus is widely recognized as the factory for ribosome assembly. In addition, recent research has revealed that the nucleolus plays essential roles in cell cycle progression, tumorigenesis, and the cellular responses to various stressors— including DNA damage (Boulon et al., 2010; Yang et al., 2018; González-Arzola, 2024). Various forms of cellular stress are accompanied by structural and functional changes in the nucleolus, underscoring its emerging function as both a sensor and regulator of stress responses. The term nucleolar stress, also known as ribosomal stress, is used to describe cellular stress that is caused by disorders of ribosome biogenesis and mostly accompanied by abnormalities of the nucleolar structure, which in turn compromise cellular homeostasis (Ohbayashi & Sugiyama, 2018). Importantly, the nucleolus is a dynamic structure in nucleus which undergoes cycles of disassembly and reassembly with every cell cycle (Dubois & Boisvert, 2016).

In animal cells, the proteins p53 and MDM2 are pivotal in mounting an effective nucleolar stress response. The transcription factor p53 not only reacts to stress signals from the nucleolus but also governs responses to other cellular stressors such as DNA damage. In contrast, plants lack the p53 and MDM2 families. Instead, members of the plant-specific NAC domain protein family have emerged as key regulators in pathways that manage various cellular stresses (Ohbayashi & Sugiyama, 2018). For instance, SUPPRESSOR OF GAMMA RESPONSE 1 (SOG1)/ANAC008 is involved in plant DNA damage responses (Preuss and Britt, 2003; Yoshiyama et al., 2009). Recent studies have indicated that the NAC domain protein SRIW1/ANAC082 participates in the plant nucleolar stress response (Ohbayashi et al., 2017; Ohbayashi & Sugiyama, 2018). However, the relationship between these pathways is still unknown.

Mutations in genes encoding nucleolar proteins in plants often result in characteristic phenotypes such as narrow, pointed leaves. Notably, double mutants involving these nucleolar protein gene defects and the *as2* mutation exhibit a marked reduction in lateral cell proliferation, leading to the formation of rod-shaped filamentous leaves. This observation strongly suggests that the nucleolus is intricately involved in key aspects of leaf development (Horiguchi et al., 2011; Matsumura et al., 2016; Ohbayashi et al., 2017).

In our study, we screened a library of natural organic compounds to identify those capable of inducing rod-shaped leaf morphology specifically within the *as2* mutant background. This screening effort yielded eight candidate molecules (Fig. 1A; Table 1). Upon further investigation, we discovered that molecules inducing the rod-shaped, filamentous leaves in the *as2* background share a common feature, they all exhibit antitumor activity in animal cells (Table 2). Notably, several of these molecules have been shown to induce nucleolar stress in our as well as other systems, hinting at a conserved mechanism of action (Boulon et al.,2010; González-Arzola 2024).

Further reinforcing the link between nucleolar stress and leaf morphology, our study shows that a mutation in the *ANAC082* gene (*sriw1*), which encodes a nucleolar stress response factor in Arabidopsis, can suppress the filamentous leaf phenotype in *as2-1 top1α-1* mutants. This finding indicates that nucleolar stress is likely induced in *top1α-1* plants. Additionally, in these plants, the typical localization of AS2 bodies within the nucleolus is disrupted, with several smaller AS2 bodies forming within the nucleolar compartment (Fig. 6B;7A). This reorganization suggests that AS2 bodies may help alleviate structural dysfunctions in the nucleolus that arise under stress.

The AS2 protein itself contains a zinc-finger motif (ZF) that binds to specific DNA sequences—an interaction that is essential for the formation of AS2 bodies and for mediating AS2 function (Vial-Pradel et al., 2018; Ando et al., 2023). Moreover, the internal-conserved G sequence (ICG) and the leucine-zipper-like sequence (LZL) within the AS2 domain are proposed to be key interacting motifs, further supporting its regulatory role (Iwakawa et al., 2002; 2020). In the context of continuous exposure to nucleolar stress during plant development, the reliable function of AS2 appears crucial for maintaining organ differentiation and ensuring stable cell proliferation.

In summary, AS2 emerges as a guardian angel in plants, protecting against the detrimental effects of nucleolar stress while facilitating proper leaf morphogenesis. By mitigating stress-induced disruptions in nucleolar function, AS2 ensures that developmental processes—such as the formation of flat leaves—proceed smoothly even under challenging conditions. This study not only underscores the multifaceted roles of the nucleolus beyond ribosome assembly but also highlights the sophisticated strategies plants employ to manage cellular stress.

Looking ahead, it will be vital to identify additional factors that interact with AS2 and further elucidate the molecular mechanisms by which it alleviates nucleolar stress. Such insights could pave the way for novel strategies to enhance plant resilience and improve crop performance under stress conditions. The interplay between stress responses, nucleolar dynamics, and developmental regulation represents a promising frontier in plant biology research.

## MATERIALS AND METHODS

### Plant strains and growth conditions

*Arabidopsis thaliana* ecotype Col-0 (CS1092) and the mutants *as1-1* (CS3374), *as2-1* (CS3117), and SALK T-DNA insertion lines were obtained from the Arabidopsis Biological Resource Center (ABRC) (Columbus, OH, USA). In this study, the T-DNA insertion lines SALK_069847, SALK_022236, SALK_089717, and SALK_069817 are designated as *top1β-1*, *top1β-2*, *krp5-4*, and *krp2-5*, respectively. The genomic maps are shown in Figs. S2A, S4A and S5A. The mutants *ett-13, bp-1 knat2-3 knat6-2*, *top1α-1*, *mgo1-7* (SALK_112625), *eal-1*, and *krp5-1* (SALK_053533) have been described previously (Takahashi et al., 2002; Graf et al., 2010; Ikezaki et al., 2010; Ishibashi et al., 2012; Iwasaki et al., 2013; Wen et al., 2013). Primers and combinations for detecting each mutation by genomic PCR are shown in Tables S5 and S6, respectively. Ages of plants are given in terms of the number of days after sowing (DAS).

### Chemicals

The library of molecules containing 2,209 natural molecules was purchased from Inter BioScreen Ltd (Moscow, Russia). 10H-CPT and etoposide were purchased from Cosmo Bio Co., Ltd. (Tokyo, Japan) and Sigma-Aldrich (St. Louis, Missouri, USA), respectively. Mitomycin C, paclitaxel, oryzalin, and propyzamide were purchased from Fujifilm Wako Pure Chemical Corporation. Bleomycin A5 was purchased from LKT Laboratories, Inc. For all chemical treatments, molecules, except mitomycin C, were dissolved in dimethyl sulfoxide (DMSO), mitomycin C was dissolved in H_2_O, and each was mixed with Murashige and Skoog (MS) agar medium and immediately dispensed into plastic multi-dishes (multidish 267061, Thermo Fisher Scientific K. K., Yokohama, Japan). 1,3,5(10)-estratriene-3,17 β-diol (17β-estradiol) was purchased from Sigma (St. Louis, MO, USA) The 17β-estradiol was prepared as a 100 μg/mL stock solution dissolved in DMSO. The solutions were stored at −20^◦^C in darkness. The working solution was made by diluting the stock solution just before use.

### Histochemical assay for GUS and GFP

Histochemical detection of GUS activity was performed essentially as described by Iwasaki et al. (2013). *BPp:GUS*/*as2-1* or 4.9 kb *ETTp:GUS*/*as2-1* (Iwasaki et al., 2013) plants were treated with and without 0.15 μM 10H-CPT for 21 days and fixed. Fixed plants were incubated in GUS staining solution described in Iwasaki et al. (2013) at 37°C for 24 h. For the preparation of *FILp:GFP*/*as2-1* plants treated with 10H-CPT or *FILp:GFP*/*as2-1 top1α-1* plants, thin sections and cleared specimens were prepared, and GFP fluorescence images were obtained with a confocal laser microscope (LSM510, Carl Zeiss Inc., Oberkochen, Germany) as described previously (Nakagawa et al., 2012).

### Plasmid construction and transformation

For preparation of *BPp:GUS*, bacterial artificial chromosome (BAC) F9M13 from chromosome IV was obtained from the *Arabidopsis* Biological Resources Center. BAC DNA was digested with *Bam*HI (at 5,3-kb upstream sequence of *BP* gene) and *NheI* (at 0.3-kb downstream of translation start site of *BP* gene), and the resultant 5,6-kb fragment of genomic DNA was inserted into pBluescript SK-, and then the product was introduced into the binary vector pGreen Km^+^ bar^+^ with the coding sequence of β-glucuronidase. We transformed the Col-0 plants using *Agrobacterium tumefaciens* GV3101 that harbored the plasmid with the incorporated DNA fragment, as described by Bechtold and Pelletier (1998). We crossed *BPp:GUS*/Col-0 plants to *as2-1* mutant plants used for our experiments.

For preparation of *Op*TOP1-expression vector(s), *OpTOP1* cDNA from the entry vectors pDONR221-OpTOP1 (Sirikantaramas et al. 2008) was transferred to the binary T-DNA destination vector pGWB2 (Nakagawa et al. 2007) in reactions mediated by LR clonase (Invitrogen, Life Technologies Japan Ltd, Tokyo, Japan). We transformed the *as1-1* plants using *Agrobacterium tumefaciens* GV3101 that harbored the resultant recombination constructs, *35Sp:OpTOP1* with the incorporated DNA fragment, as described by Bechtold and Pelletier (1998).

For construction of *CYCB1;2p:GUS,* the promoter fragment (1.0 kb) from *CYCB1;2* gene was cloned into the entry vector pENTR-D/TOPO (ThermoFisher), and the resulting plasmid was subjected to LR reaction with the destination vectors pBGGUS (Kubo et al., 2005) to create *CYCB1;2p:GUS*. The construct was introduced by floral-dip transformation (Clough and Bent, 1998) into Arabidopsis (Col-0) plants. For cloning of the promoter fragment from *CYCB1;2* gene locus, the primers Cyc1b fullpro5/CACC: CACCATCGTGAAGGTAACATTTACAAC and Cyc1b-p3: TTCTCTTTCGTAAAGAGTCTCTGCG were used.

### Quantification of GUS activity

The quantitative GUS assay was performed as described previously (Halder and Kombrink 2015; Iwakawa et al. 2021), with some modifications. Aerial part from 13-day-old *CYCB1;2p:GUS* seedlings grown in 1/2MS medium were harvested in zirconia beads-containing 2 ml tubes, crushed with Tissue lyzer, resuspended with 200 μL of extraction buffer containing 100 mM sodium phosphate pH 7.0 and 1 mM dithiothreitol, and centrifuged at 15,000 rpm at 4◦C for 20 min. After determining the protein concentrations of the supernatants, 160 μL of the supernatant (corresponding to 30 μg protein) was transferred to a 96-well transparent plate and mixed with GUS assay solution (40 μL) containing 5 mM 4-methylumbelliferyl-β-D-glucuronide (TCI M3026), 125 mM sodium phosphate pH 7.0, 2.5 mM EDTA, 0.25% Triton X-100, and 25 mM β-mercaptoethanol. Samples were incubated at 37◦C for 60 min, and 4-methylumbelliferone (4-MU) fluorescence was measured every 10 min using the plate reader Spark (TECAN, Switzerland) with an excitation/emission wavelength of 365/455 nm. The GUS activity was calculated using theΔE455 increments (10–120 min).

### Real time quantitative reverse transcriptase-polymerase chain reaction (qRT-PCR)

Each 30 shoot apices from 14-day-old plants were collected, and total RNA was extracted with the RNeasy Plant Mini Kit (Qiagen, Valencia, CA) according to the manufacturer’s instructions. Sample volumes were normalized as described by Nakagawa et al. (2012). The primer pairs are shown in Table S7. For the analysis of RNA levels in Arabidopsis by real-time qRT-PCR, we prepared 10 µg of total RNA. Reverse transcription was carried out with ReverTra Ace (TOYOBO, Osaka, Japan). PCR was performed in the presence of the double-stranded DNA-specific dye SYBR Green (Applied Biosystems, Lincoln, CA). Amplification was monitored in real time with the Applied Biosystems StepOnePlus Real-Time PCR system (Applied Biosystems) according to the supplier’s recommendations. The mean values of three technical replicates were normalized by using the *ACTIN2* (*ACT2*) or *TRANSLATION ELONGATION FACTOR* (*EF-1α*) transcripts as each control.

### Introduction of AS2-YFP into *top1α-1*

We crossed the Col-0 background AS2-YFP stamens with *top1α-1* pistils. The F_1_ seeds were selected on MS medium that contained hygromycin. Then, the F_2_ plants were selected on MS medium supplemented with hygromycin, based on leaf phenotype, via PCR. The appropriate pairs of primers for each mutant that we used for the genotyping are listed in Table S7. We used the F_3_ plants for the AS2-YFP observation. To examine the subcellular localizations of AS2-YFP, images were recorded via confocal laser scanning fluorescence microscopy (LSM710; Carl Zeiss, AG, Baden-Württemberg, Germany). We acquired z-stack images of the AS2 bodies using Zen 2012 software (Carl Zeiss AG, Baden-Württemberg, Germany) as described previously (Ando et al., 2023).

### Observation of Fluorescence

To examine the subcellular localizations of the YFP fused to AS2, we established three lines of transgenic Arabidopsis plants. To induce the expression of the transgenes, the 6 DAS transgenic Arabidopsis whole plants were soaked in 0.05 μg/mL of 17β-estradiol in 1× MS medium under vacuum conditions for 30 min, and were then incubated at 22 ^◦^C overnight. The patterns of fluorescence due to YFP were analyzed in 4 to 75 cells of each line of transgenic plants after the induction of expression of transgenes. YFP-positive nuclei were observed in the adaxial epidermis of the cotyledons. Images were recorded via confocal laser scanning fluorescence microscopy with an Objective Plan-Apochromat 63×/1.4 Oil DICII and an Objective α Plan-Apochromat 100×/1.46 Oil DIC (LSM710; Carl Zeiss, AG, Baden-Württemberg, Germany). We acquired z-stack images of the AS2 bodies using Zen 2012 software (Carl Zeiss AG, Baden-Württemberg, Germany).

### Microarray

Shoot apices from seedlings of wild-type and *as2-1* plants grown with and without 0.15 µM 10H-CPT for 14 days were harvested and total RNA was extracted as described in the ‘real time qRT-PCR’ section, above. For microarray analysis, the quality and purity of the RNA were confirmed with an Ultrospec 2100 pro (GE Healthcare UK, Ltd, Amersham Place, England). Total RNA samples (100 ng) were reverse-transcribed, yielding double-stranded cDNA, which was transcribed *in vitro* in the presence of biotin-labeled nucleotides with an IVT Labeling Kit (Affymetrix Inc., Santa Clara, CA), and purified. Labeled cRNA was fragmented and hybridized to Affymetrix ATH1 GeneChip arrays for 16 hours at 45°C according to Affymetrix protocols. Arrays were washed on an Affymetrix Fluidics Station 450 and measured for fluorescence intensity with an Affymetrix GeneChip Scanner 3000. The raw data were processed by using Affymetrix Gene Chip Operating Software (GCOS; Version 1.4.0.036).

### MM2d cells and synchronization

A suspension culture of *Arabidopsis thaliana* cell line MM2d (L*er* ecotype) (Menges and Murray, 2002) was maintained at 25°C in the dark by weekly subculturing in modified Linsmaier and Skoog medium (Haga et al., 2007) with rotation at 120 rpm. To synchronize the MM2d cell cycle at the G1/S phase boundary, 40 ml of a 7-day-old cell culture were transferred to 160 ml of modified Linsmaier and Skoog medium containing 5 mg/L aphidicolin (Wako Pure Chemicals). After culturing for 24 hours, cells were washed and cultured further in 200 ml of fresh medium. At every 2 hours after the drug removal, cells were harvested by centrifugation, frozen in liquid nitrogen and stored at –80°C until use. RNA was extracted with Trizol reagent according to the manufacturer’s instructions (Invitrogen). Purification of Poly(A)^+^ RNA and synthesis of first-strand cDNA were performed as described previously (Iwakawa et al. 2007).

## Supporting information

2_20250922_Nakagawa-et_al_Supplementary information

## Acknowledgements

We are grateful to Dr. Taku Takahashi (Okayama University), Dr. Tsuyoshi Nakagawa (Shimane University), and Dr. Kazuki Saito (Chiba University) for providing *top1α-1* seeds, the pGWB2 plasmid, and *OpTOP1* cDNA, respectively. We thank Messrs. Fumiaki Ogasawara (Nagoya University), Takao Yamamoto, Toshiki Ohkochi, Kazuaki Kawasaki (Chubu University), Mses. Mayu Sakuma, Yuka Atsumi, Mari Morimoto, and Misato Yamakawa (Chubu University) for technical assistance. We are grateful to Dr. Kazuki Saito (Chiba University), Dr. Taisuke Nishimura (Nagaoka University of Technology), Dr. Yuichiro Tsuchiya, Dr. Ayato Sato and Dr. Yoko Matsumura (Nagoya University), Dr. Johji Miwa (IBR) for helpful discussion and cooperation.

## Author contributions

Conceptualization: A.N., H.I., H.T., Y.M., C.M.;Methodology: A.N., H.I., H.T., C.M.;Validation: A.N., H.I., H.T., C.M.;Formal analysis: A.N., H.I., H.T., S.V-P., Ma.T., K.O., T.I., Mo.T., E.T., N.I., S.A.; Investigation: A.N., Su.K., Sh.K., M.S. C.M.;Resources: M.I., M.Y., B.-Y.C., J.-T.W., I.O., Mu.S.;Data curation: A.N., H.I., H.T.;Writing - original draft: A.N., H.I., H.T., C.M.;Writing - review & editing: A.N., H.I., IO., Sh.K., Mu.S., Mi.S., Y.M., C.M.;Visualization: A.N., H.I., C.M.;Supervision: C.M.; Project administration: Y.M., C.M.;Funding acquisition: Sh.K., Mi.S., Y.M., C.M.,

## Data availability

All relevant data can be found within the article and its supplementary information. Original microarray data were deposited in DDBJ GEA under accession number ESUB002266.

## Supplementary information

**Supplementary Figure S1. Mutations of class 1 *KNOX* genes did not suppress the phenotype of filamentous leaves.**

**Supplementary Figure S2. Mutation of *TOP1α*/*MGOUN1* (*MGO1*) enhanced the leaf abaxialization of *as1* and *as2,* but mutation of *TOP1β* did not.**

**Supplementary Figure S3. The microarray and KB-FuzzyART classified 30 clusters and the *KRP5*/*ICK3* gene is the only gene that is categorized as a cell division gene.**

**Supplementary Figure S4. The *KRP5*/*ICK3* gene is responsible for the formation of filamentous leaves in 10H-CPT-treated *as2*.**

**Supplementary Figure S5. The *KRP2*/*ICK2* gene was not responsible for the formation of filamentous leaves in *top1α as2* and 10H-CPT-treated *as2*.**

**Supplementary Figure S6. Inhibitors of DNA replication induce formation of filamentous leaves, but treatment with oryzalin and paclitaxel did not exhibit defects in leaf adaxial/abaxial polarity.**

**Table S1. The constructed clusters and assignment of genes for Dataset-C**

**Table S2. Enrichment rate of each gene cluster for Gene list-4.**

**Table S3. *p* values for enrichment rate of each gene cluster and each category in Gene list-4.**

**Table S4. *q* value for enrichment rate of each gene cluster and each category in Gene list-4.**

**Table S5. Sequences for primers used for genomic PCR.**

**Table S6. Combinations of primers for detection of mutation or T-DNA insertion by genomic PCR.**

**Table S7. Sequences for primers used for real-time RT-PCR.**

