## Supplementary material for "Arabidopsis epigenetic factor AS2 attenuates nucleolar stress by camptothecin and establishes leaf polarity by repressing a CDK inhibitor": 2_20250922_Nakagawa-et_al_Supplementary information: 2_20250922_Nakagawa-et_al_Supplementary information.pdf

**6 Supplementary Figures**

**7 Supplementary Tables**

#### Figure S1

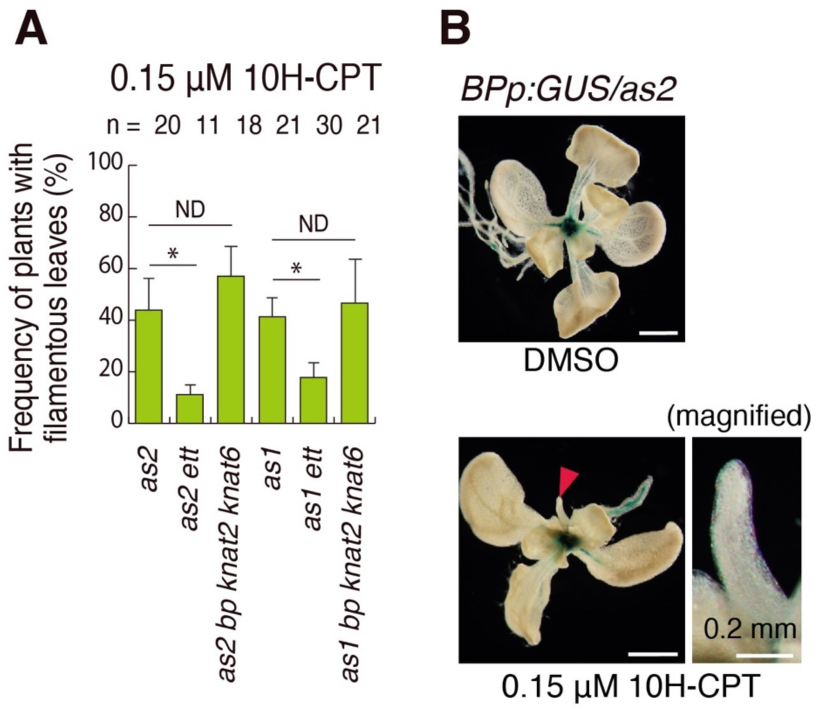


**Supplementary Figure S1. Mutations of class 1 *KNOX* genes did not suppress the phenotype of filamentous leaves.**

(A) Frequencies of plants with filamentous leaves at positions of 1st and 2nd leaves when plants were treated with 0.15 µM 10H-CPT. ($p$<0.05, Tukey's test after significant ANOVA). (B) A typical *BPp:GUS*/*as2-1* plant treated with 0.15 µM 10H-CPT for 14 days. Bars = 1 mm.

#### Figure S2

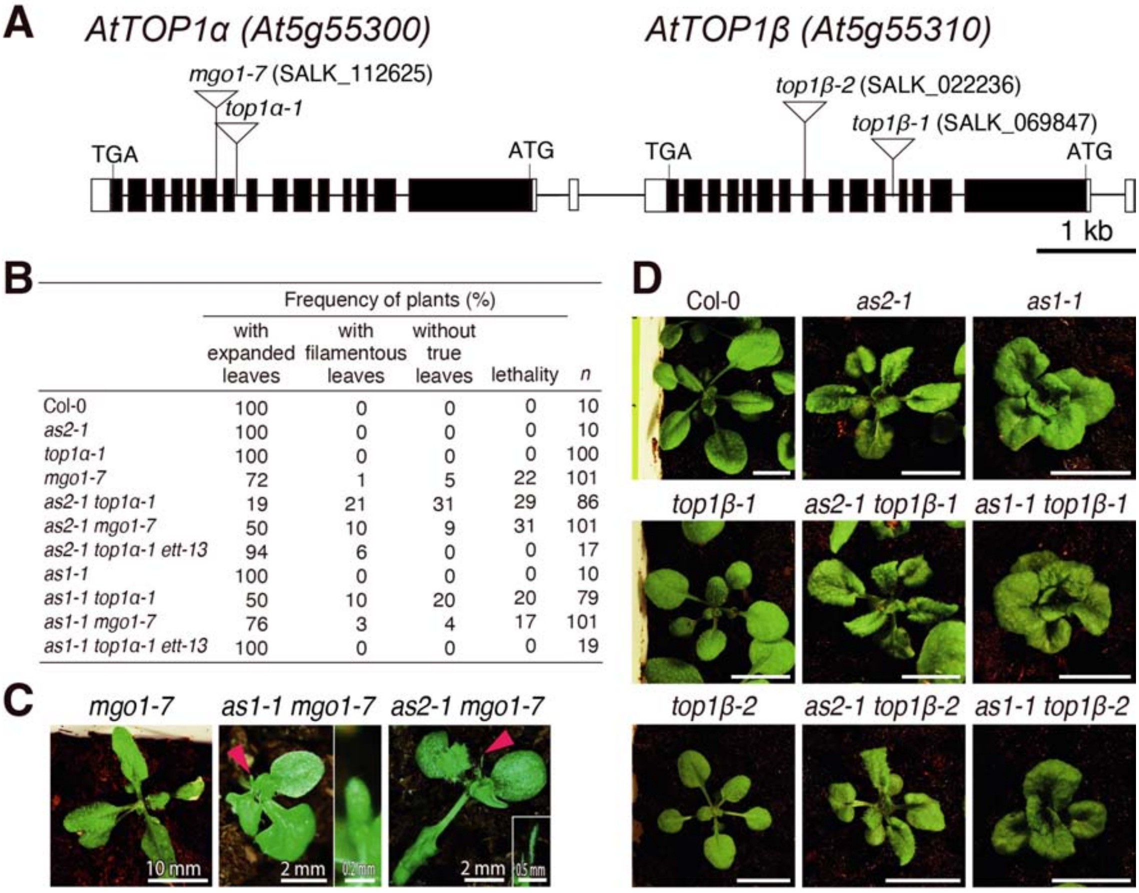


| | Frequency of plants (%) | | | | |
|---|---|---|---|---|---|
| | with expanded leaves | with filamentous leaves | without true leaves | lethality | *n* |
| Col-0 | 100 | 0 | 0 | 0 | 10 |
| *as2-1* | 100 | 0 | 0 | 0 | 10 |
| *top1α-1* | 100 | 0 | 0 | 0 | 100 |
| *mgo1-7* | 72 | 1 | 5 | 22 | 101 |
| *as2-1 top1α-1* | 19 | 21 | 31 | 29 | 86 |
| *as2-1 mgo1-7* | 50 | 10 | 9 | 31 | 101 |
| *as2-1 top1α-1 ett-13* | 94 | 6 | 0 | 0 | 17 |
| *as1-1* | 100 | 0 | 0 | 0 | 10 |
| *as1-1 top1α-1* | 50 | 10 | 20 | 20 | 79 |
| *as1-1 mgo1-7* | 76 | 3 | 4 | 17 | 101 |
| *as1-1 top1α-1 ett-13* | 100 | 0 | 0 | 0 | 19 |

**Supplementary Figure S2. Mutation of *TOP1α/MGOUN1* (*MGO1*) enhanced the leaf abaxialization of *as1* and *as2,* but mutation of *TOP1β* did not.**

(A) Genomic structures (exon-intron organizations) of type IB DNA topoisomerases, *AtTOP1α/MGO1* and *AtTOP1β*. Black boxes indicate exons, and white boxes indicate untranslated exons. Sites of T-DNA insertions in mutants used in this study are indicated. (B) The *top1α-1/mgo1-7* mutation in *as2* or *as1* background exhibited enhanced formation of filamentous/trumpet-shaped leaves. Filamentous leaves and other abnormal shapes of leaves such as trumpet-shaped leaves were counted in the indicated genetic backgrounds. The *ett-13* mutation suppressed the filamentous leaves phenotype. (C) The mutation of *mgo1-7* exhibited impairment of leaf adaxial/abaxial polarity in *as1* and *as2* backgrounds. The phenotypes of plants grown in soil for 21 days are shown. Red arrowheads indicate filamentous leaves. (D) Genetic interactions between *AS1*, *AS2*, and *AtTOP1β*. The phenotypes of plants grown in soil for 21 or 22 days are shown. Bars = 10 mm.

#### Figure S3

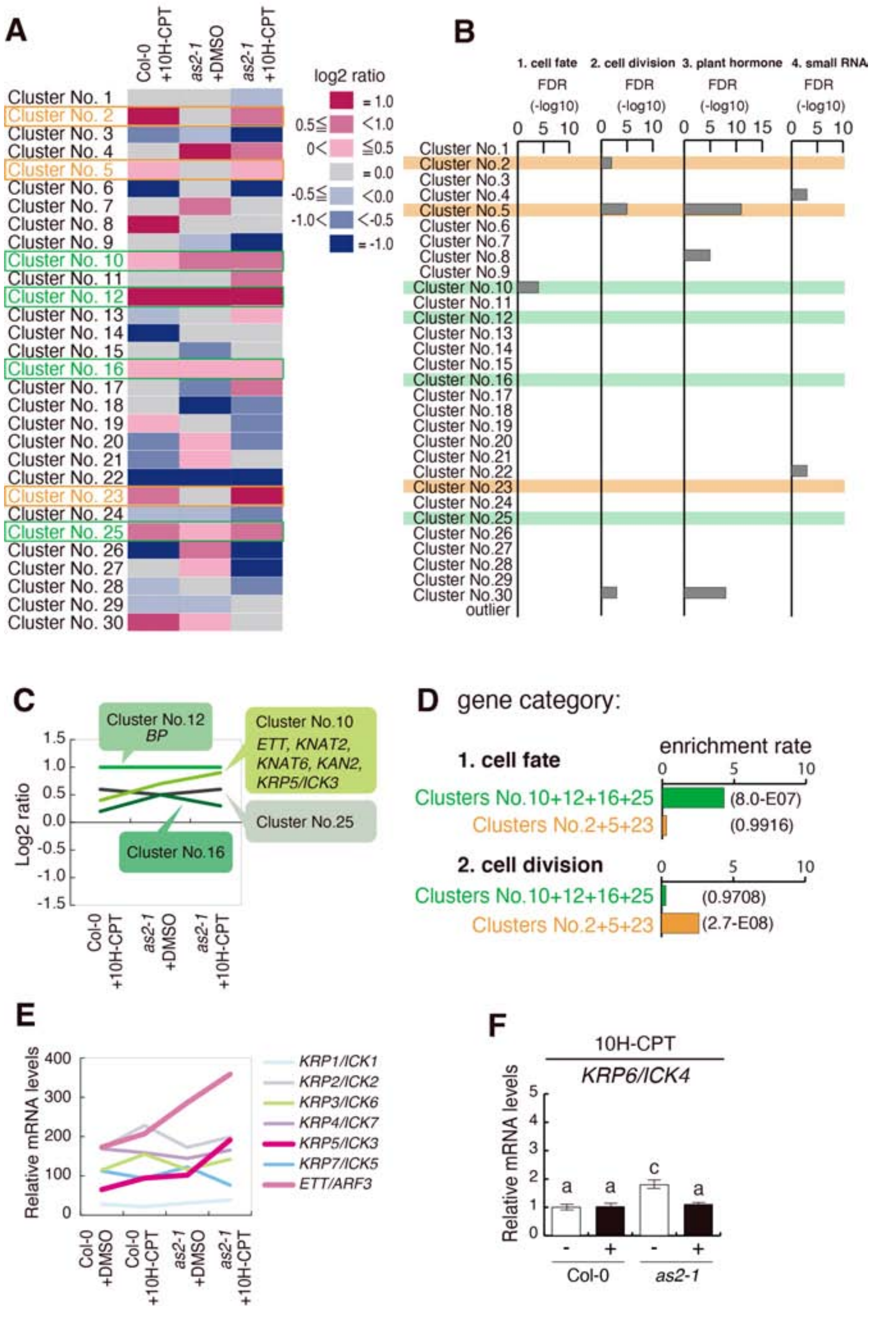


**Supplementary Figure S3. The microarray and KB-FuzzyART classified 30 clusters and the *KRP5*/*ICK3* gene is the only gene that is categorized as a cell division gene.**

(A) Clustering analysis by KB-FuzzyART. Levels of gene expression in comparison with those of Col-0 treated with DMSO are shown. (B) Gene enrichment analysis. Enrichment rates for 2 categories (cell fate and cell division) in gene list-4 among 8,009 genes were calculated. Values indicate –log10 q. (C) The expression patterns for cluster that includes *ETT* (cluster 10) and those for similar expression patterns (clusters 12, 16, and 25). (D) Gene enrichment analysis of clusters 10 + 12 + 16 + 25 and 2 + 5 + 23. Enrichment rates were calculated for 2 categories (cell fate and cell division) in gene list-4 among 8,009 genes. Numbers in parenthesis indicate q values. (E) The transcript levels of *ETT*/*ARF3* and *KRP*/*ICK3* genes (except *KRP6*/*ICK4*) in shoot apices of plants treated with and without 10H-CPT. (F) The transcript levels of *KRP6*/*ICK4* in shoot apices of plants treated with and without 10H-CPT ($p$<0.05, Tukey's HSD test after significant ANOVA).

### Figure S4

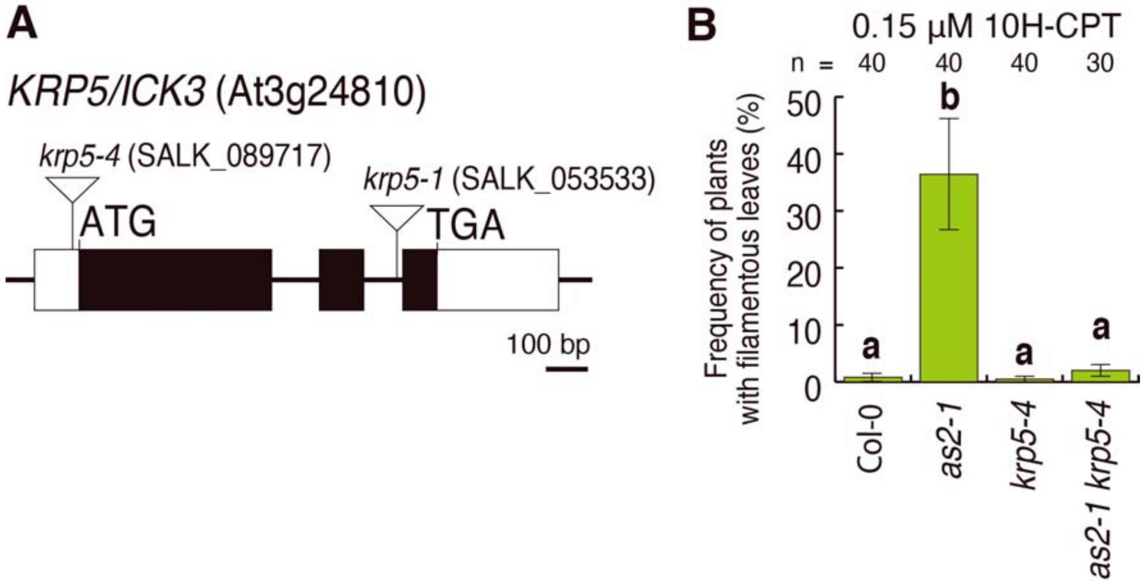


**Supplementary Figure S4. The *KRP5*/*ICK3* gene is responsible for the formation of filamentous leaves in 10H-CPT-treated *as2*.**

(A) Genomic structures of *KRP5*/*ICK3*. Black boxes indicate exons, and white boxes indicate untranslated exons. Sites of T-DNA insertions of mutants used in this study are indicated. (B) Frequencies of *krp5-4* plants with filamentous leaves when plants were treated with 0.15 µM 10H-CPT. Data represent mean ±SD of 3 replicates (n=40 or 30 for each).

### Figure S5

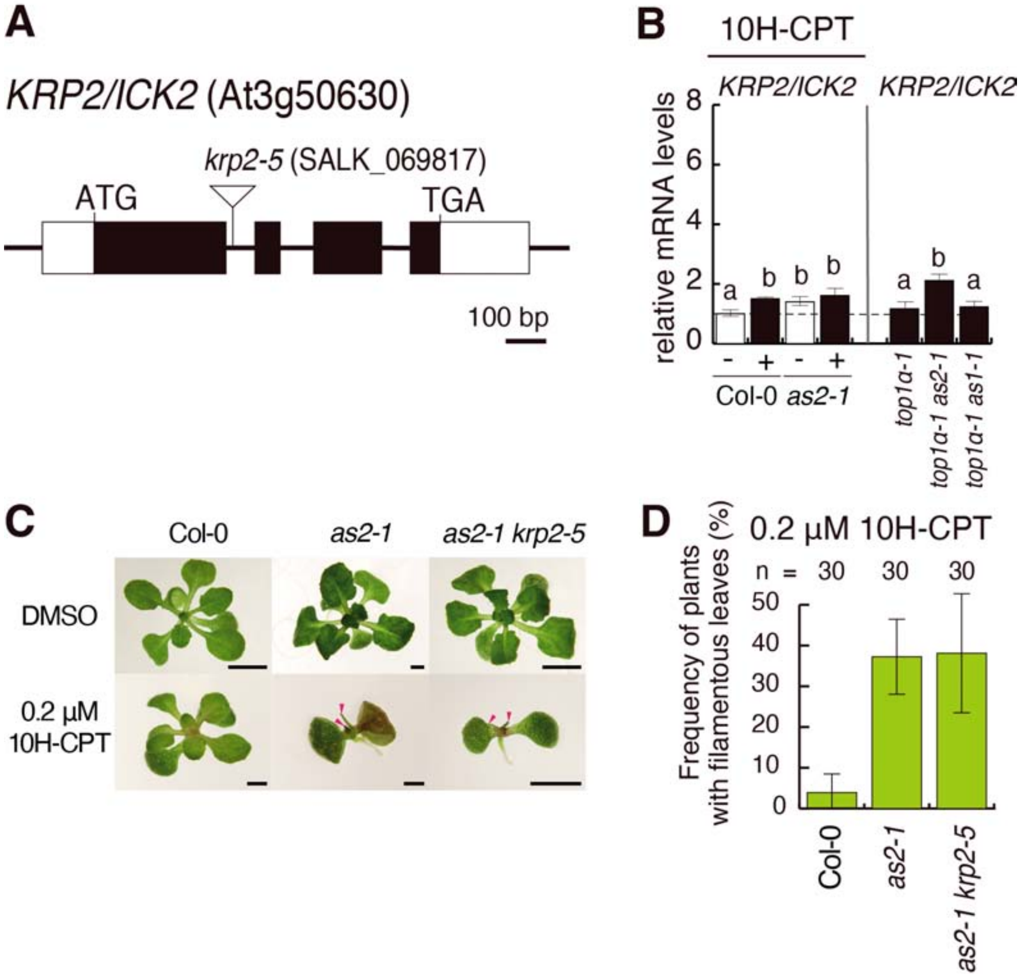


**Supplementary Figure S5. The *KRP2*/*ICK2* gene was not responsible for the formation of filamentous leaves in *top1α as2* and 10H-CPT-treated *as2*.**

(A) Genomic structures of *KRP2*/*ICK2*. Black boxes indicate exons, and white boxes indicate untranslated exons. Sites of T-DNA insertions of mutants used in this study are indicated. (B) Changes in transcription levels of *KRP2*/*ICK2*. Letters above bars indicate significance among transcript of each gene ($p$<0.05, Tukey's HSD test after significant ANOVA). (C) Phenotypes of plants treated with 0.2 µM 10H-CPT for 14 days. Bars = 1 mm. Red arrowheads indicate filamentous leaves. (D) Frequencies of plants with filamentous leaves when plants were treated with 0.2 µM 10H-CPT. Data represent mean ±SD of 3 replicates (n=30 for each).

#### Figure S6

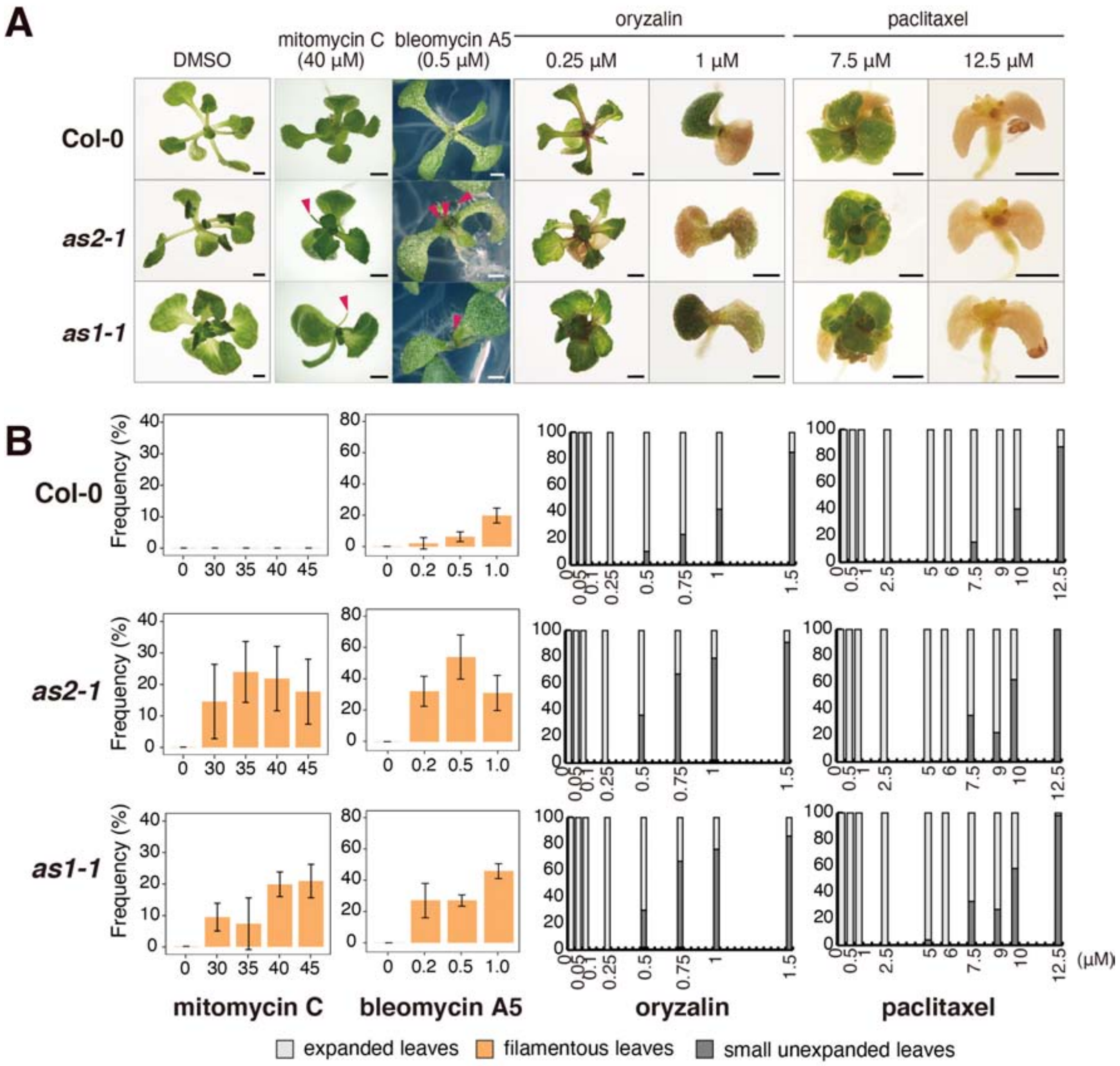


**Supplementary Figure S6. Inhibitors of DNA replication induce formation of filamentous leaves, but treatment with oryzalin and paclitaxel did not exhibit defects in leaf adaxial/abaxial polarity.**

(A) Representative phenotypes of plants treated with mitomycin C, bleomycin A5, oryzalin, or paclitaxel for 21 days. Red arrowheads indicate filamentous leaves. Bars = 1 mm. (B) Dose-dependent effects of mitomycin C, bleomycin A5 oryzalin and paclitaxel on leaf abnormality and leaf formation. Each value is calculated from the means of 3 biological replicates (n=20 for each).

**Table S1.** The constructed clusters and assignment of genes for Dataset-C

| Cluster number | No. of genes | No. of genes in Gene list-4 | Categories in Gene list-4 | | | | Pattern | | |
|---|---|---|---|---|---|---|---|---|---|
| | | | 1. cell fate | 2. cell division | 3. plant hormone | 4. small RNA | Col-0 10H-CPT | *as2-1* DMSO | *as2-1* 10H-CPT |
| 1 | 956 | 15 | 5 | 5 | 3 | 2 | 0 | 0 | - |
| 2 | 205 | 7 | 0 | 6 | 1 | 0 | + | 0 | + |
| 3 | 160 | 7 | 2 | 0 | 5 | 0 | - | - | - |
| 4 | 145 | 10 | 2 | 0 | 8 | 0 | 0 | + | + |
| 5 | 1507 | 35 | 3 | 24 | 7 | 1 | + | 0 | + |
| 6 | 267 | 8 | 1 | 0 | 7 | 0 | - | 0 | - |
| 7 | 246 | 10 | 2 | 0 | 7 | 1 | 0 | + | 0 |
| 8 | 151 | 4 | 0 | 3 | 1 | 0 | + | 0 | 0 |
| 9 | 181 | 10 | 4 | 1 | 4 | 1 | 0 | - | - |
| 10 | 140 | 9 | 7 | 1 | 1 | 0 | + | + | + |
| 11 | 980 | 13 | 3 | 0 | 8 | 2 | 0 | 0 | + |
| 12 | 170 | 5 | 2 | 0 | 3 | 0 | + | + | + |
| 13 | 364 | 3 | 1 | 0 | 2 | 0 | - | 0 | + |
| 14 | 287 | 3 | 0 | 0 | 3 | 0 | - | 0 | 0 |
| 15 | 317 | 7 | 1 | 2 | 4 | 0 | 0 | - | 0 |
| 16 | 174 | 9 | 5 | 0 | 2 | 2 | + | + | + |
| 17 | 19 | 1 | 0 | 0 | 1 | 0 | 0 | - | + |
| 18 | 71 | 4 | 0 | 0 | 4 | 0 | 0 | - | - |
| 19 | 34 | 1 | 0 | 0 | 1 | 0 | + | 0 | - |
| 20 | 30 | 1 | 1 | 0 | 0 | 0 | - | + | - |
| 21 | 70 | 1 | 0 | 0 | 1 | 0 | - | + | 0 |
| 22 | 181 | 9 | 0 | 0 | 9 | 0 | - | - | - |
| 23 | 204 | 4 | 1 | 0 | 3 | 0 | + | 0 | + |
| 24 | 94 | 3 | 1 | 0 | 2 | 0 | - | - | - |
| 25 | 72 | 1 | 1 | 0 | 0 | 0 | + | + | + |
| 26 | 22 | 1 | 1 | 0 | 0 | 0 | - | + | - |
| 27 | 136 | 7 | 4 | 0 | 3 | 0 | 0 | + | - |
| 28 | 373 | 4 | 1 | 1 | 2 | 0 | - | 0 | - |
| 29 | 173 | 1 | 1 | 0 | 0 | 0 | - | - | 0 |
| 30 | 120 | 7 | 1 | 6 | 0 | 0 | + | + | 0 |
| Outlier | 160 | 0 | 0 | 0 | 0 | 0 | | | |
| Over all | 8009 | 200 | 50 | 49 | 92 | 9 | | | |

+, -, and 0 indicate up-regulation, down-regulation, and no change, respectively.

**Table S2.** Enrichment rate of each gene cluster for Gene list-4.

| Cluster No. | Categories in Gene list-4 | | | |
|---|---|---|---|---|
| | 1. cell fate | 2. cell division | 3 plant hormone | 4. small RNA |
| 1 | 0.8 | 0.9 | 0.3 | 1.9 |
| 2 | 0.0 | 4.8 | 0.4 | 0.0 |
| 3 | 2.0 | 0.0 | 2.7 | 0.0 |
| 4 | 2.2 | 0.0 | 4.8 | 0.0 |
| 5 | 0.3 | 2.6 | 0.4 | 0.6 |
| 6 | 0.6 | 0.0 | 2.3 | 0.0 |
| 7 | 1.3 | 0.0 | 2.5 | 3.6 |
| 8 | 0.0 | 3.2 | 0.6 | 0.0 |
| 9 | 3.5 | 0.9 | 1.9 | 4.9 |
| 10 | 8.0 | 1.2 | 0.6 | 0.0 |
| 11 | 0.5 | 0.0 | 0.7 | 1.8 |
| 12 | 1.9 | 0.0 | 1.5 | 0.0 |
| 13 | 0.4 | 0.0 | 0.5 | 0.0 |
| 14 | 0.0 | 0.0 | 0.9 | 0.0 |
| 15 | 0.5 | 1.0 | 1.1 | 0.0 |
| 16 | 4.6 | 0.0 | 1.0 | 10.2 |
| 17 | 0.0 | 0.0 | 4.6 | 0.0 |
| 18 | 0.0 | 0.0 | 4.9 | 0.0 |
| 19 | 0.0 | 0.0 | 2.6 | 0.0 |
| 20 | 5.3 | 0.0 | 0.0 | 0.0 |
| 21 | 0.0 | 0.0 | 1.2 | 0.0 |
| 22 | 0.0 | 0.0 | 4.3 | 0.0 |
| 23 | 0.8 | 0.0 | 1.3 | 0.0 |
| 24 | 1.7 | 0.0 | 1.9 | 0.0 |
| 25 | 2.2 | 0.0 | 0.0 | 0.0 |
| 26 | 7.3 | 0.0 | 0.0 | 0.0 |
| 27 | 4.7 | 0.0 | 1.9 | 0.0 |
| 28 | 0.4 | 0.4 | 0.5 | 0.0 |
| 29 | 0.9 | 0.0 | 0.0 | 0.0 |
| 30 | 1.3 | 8.2 | 0.0 | 0.0 |
| Outlier | 0.0 | 0.0 | 0.0 | 0.0 |
| 02+05+23 | 0.3 | 2.6 | 0.5 | 0.5 |
| 10+12+16+25 | 4.3 | 0.3 | 0.9 | 3.2 |
| 8+30 | 0.6 | 5.4 | 0.3 | 0.0 |

**Table S3.** *p* values for enrichment rate of each gene cluster and each category in Gene list-4.

| Cluster No. | Categories in Gene list-4 | | | |
|---|---|---|---|---|
| | 1. cell fate | 2. cell division | 3 plant hormone | 4. small RNA |
| 1 | 0.73 | 0.71 | 1.00 | 0.29 |
| 2 | 1.00 | 1.5E-03 | 0.91 | 1.00 |
| 3 | 0.26 | 1.00 | 3.7E-02 | 1.00 |
| 4 | 0.23 | 1.00 | 2.5E-04 | 1.00 |
| 5 | 1.00 | 1.6E-06 | 1.00 | 0.85 |
| 6 | 0.82 | 1.00 | 3.3E-02 | 1.00 |
| 7 | 0.46 | 1.00 | 2.3E-02 | 0.24 |
| 8 | 1.00 | 0.06 | 0.83 | 1.00 |
| 9 | 2.6E-02 | 0.67 | 0.15 | 0.19 |
| 10 | 2.3E-05 | 0.58 | 0.80 | 1.00 |
| 11 | 0.95 | 1.00 | 0.89 | 0.30 |
| 12 | 0.29 | 1.00 | 0.31 | 1.00 |
| 13 | 0.90 | 1.00 | 0.93 | 1.00 |
| 14 | 1.00 | 1.00 | 0.65 | 1.00 |
| 15 | 0.87 | 0.58 | 0.50 | 1.00 |
| 16 | 4.4E-03 | 1.00 | 0.60 | 1.5E-02 |
| 17 | 1.00 | 1.00 | 0.20 | 1.00 |
| 18 | 1.00 | 1.00 | 8.8E-03 | 1.00 |
| 19 | 1.00 | 1.00 | 0.33 | 1.00 |
| 20 | 0.17 | 1.00 | 1.00 | 1.00 |
| 21 | 1.00 | 1.00 | 0.56 | 1.00 |
| 22 | 1.00 | 1.00 | 2.2E-04 | 1.00 |
| 23 | 0.73 | 1.00 | 0.42 | 1.00 |
| 24 | 0.45 | 1.00 | 0.29 | 1.00 |
| 25 | 0.36 | 1.00 | 1.00 | 1.00 |
| 26 | 0.13 | 1.00 | 1.00 | 1.00 |
| 27 | 1.0E-02 | 1.00 | 0.21 | 1.00 |
| 28 | 0.91 | 0.90 | 0.93 | 1.00 |
| 29 | 0.67 | 1.00 | 1.00 | 1.00 |
| 30 | 0.53 | 8.2E-05 | 1.00 | 1.00 |
| Outlier | 1.00 | 1.00 | 1.00 | 1.00 |
| 02+05+23 | 1.00 | 2.7E-08 | 1.00 | 0.91 |
| 10+12+16+25 | 8.0E-07 | 0.97 | 0.62 | 0.13 |
| 8+30 | 0.82 | 3.2E-05 | 0.96 | 1.00 |

**Table S4.** *q* value for enrichment rate of each gene cluster and each category in Gene list-4.

| Cluster No. | Categories in Gene list-4 | | | |
|---|---|---|---|---|
| | 1. cell fate | 2. cell division | 3 plant hormone | 4. small RNA |
| 1 | 1.00 | 1.00 | 1.00 | 1.00 |
| 2 | 1.00 | 1.5E-02 | 1.00 | 1.00 |
| 3 | 0.99 | 1.00 | 0.19 | 1.00 |
| 4 | 0.99 | 1.00 | 3.8E-03 | 1.00 |
| 5 | 1.00 | 4.92E-05 | 1.00 | 1.00 |
| 6 | 1.00 | 1.00 | 0.19 | 1.00 |
| 7 | 1.00 | 1.00 | 0.18 | 1.00 |
| 8 | 1.00 | 0.50 | 1.00 | 1.00 |
| 9 | 0.20 | 1.00 | 0.68 | 1.00 |
| 10 | 7.09E-04 | 1.00 | 1.00 | 1.00 |
| 11 | 1.00 | 1.00 | 1.00 | 1.00 |
| 12 | 0.99 | 1.00 | 0.84 | 1.00 |
| 13 | 1.00 | 1.00 | 1.00 | 1.00 |
| 14 | 1.00 | 1.00 | 1.00 | 1.00 |
| 15 | 1.00 | 1.00 | 1.00 | 1.00 |
| 16 | 0.07 | 1.00 | 1.00 | 0.47 |
| 17 | 1.00 | 1.00 | 0.71 | 1.00 |
| 18 | 1.00 | 1.00 | 0.09 | 1.00 |
| 19 | 1.00 | 1.00 | 0.84 | 1.00 |
| 20 | 0.89 | 1.00 | 1.00 | 1.00 |
| 21 | 1.00 | 1.00 | 1.00 | 1.00 |
| 22 | 1.00 | 1.00 | 3.8E-03 | 1.00 |
| 23 | 1.00 | 1.00 | 0.99 | 1.00 |
| 24 | 1.00 | 1.00 | 0.84 | 1.00 |
| 25 | 1.00 | 1.00 | 1.00 | 1.00 |
| 26 | 0.80 | 1.00 | 1.00 | 1.00 |
| 27 | 0.10 | 1.00 | 0.71 | 1.00 |
| 28 | 1.00 | 1.00 | 1.00 | 1.00 |
| 29 | 1.00 | 1.00 | 1.00 | 1.00 |
| 30 | 1.00 | 1.28E-03 | 1.00 | 1.00 |
| Outlier | 1.00 | 1.00 | 1.00 | 1.00 |

**Table S5**. Sequences for primers used for genomic PCR.

| Gene | Primer name | Sequence |
|---|---|---|
| *TOP1α* | TOP1α 2711 FW | 5'-ATCATGAGGCCGACACTGTT-3' |
| | TOP1α 2948 RE | 5'-TGTCCAATGGCCTTGTAAAC-3' |
| | TOP1α 2520 FW | 5'-AAATTGCTGTGGCGACATATC-3' |
| | TOP1α 3494 RE | 5'-ATTGCAGATTATGGCAACCTG-3' |
| | top1α LB | 5'-ACCCCAGTACATTAAAAACGTC-3' |
| *TOP1β* | TOP1β +1292 FW | 5'-TTCCCAGAGTGTTTTTGCATC-3' |
| | TOP1β +2409 RE | 5'-GGTAAGAAATGGGAAAGCAGG-3' |
| | TOP1β +2409 FW | 5'-CCTGCTTTCCCATTTCTTACC-3' |
| | TOP1β +3560 RE | 5'-CTCCGTGAGATTTCGAGACAG-3' |
| *ETT* | ETT LP2 | 5'-TCTTCTTTTGGCTGAGAGGG-3' |
| | ETT RP3 | 5'-GAGATAAGGGTCCAGCACA-3' |
| *pGWB2/OpTOP1* | pGWB2 35Sp +5682FW | 5'-GACGTTCCAACCACGTCTTC-3' |
| | OpTOP1 148RE | 5’-GTATGCACAGCAGATTGTCC-3’ |
| *KRP5* | KRP5 -196FW | 5'-GATGACAATCAGGTGAGAGG-3' |
| | KRP5 +73RE | 5'-GCTCTGGTACGGAAACCTAGC-3' |
| *KRP2* | KRP2 +202FW | 5'-TACGACGGCGAGATTCTCCT-3' |
| | KRP2 +440RE | 5'-CACGACGTTTCTGTTTCACG-3' |
| SALK LB | LBb1 | 5'-GCGTGGACCGCTTGCTGCAACT-3' |
| | LBb1.3 | 5’-ATTTTGCCGATTTCGGAAC-3’ |

**Table S6.** Combinations of primers for detection of mutation or T-DNA insertion by genomic PCR.

| allele or transgenic lines | | FW | RE | DNA size (bp) |
|---|---|---|---|---|
| *top1α-1* | wild type | TOP1α 2711 FW | TOP1α 2948 RE | 258 |
| | mutant | TOP1α 2948 RE | top1α LB | approx.200 |
| *mgo1-7* (SALK_112625) | wild type | TOP1α 2520 FW | TOP1α 3494 RE | 994 |
| | mutant | TOP1α 2520 FW | LBb1 | 646 |
| *top1β-1* (SALK_069847) | wild type | TOP1β +1292 FW | TOP1β +2409 RE | 1138 |
| | mutant | TOP1β +2409 RE | LBb1 | 600 |
| *top1β-2* (SALK_022236) | wild type | TOP1β +2409 FW | TOP1β +3560 RE | 1153 |
| | mutant | TOP1β +3560 RE | LBb1 | approx.700 |
| *ett-13* | wild type | ETT LP2 | ETT RP3 | 300 |
| | mutant | ETT LP2 | LBb1.3 | 367 |
| *OpTOP1* transgenic plants | T-DNA insertion | pGWB2 35Sp +5682FW | OpTOP1 148RE | 448 |
| *krp5-1* (SALK_053533) | wild type | KRP5 p2f | KRP5 p2r | 159 |
| | mutant | LBb1.3 | KRP5 p2r | 353 |
| *krp5-4* (SALK_089717) | wild type | KRP5 -196 FW | KRP5 +73 RE | 269 |
| | mutant | LBb1.3 | KRP5 +73 RE | 392 |
| *krp2-5* (SALK_069817) | wild type | KRP2 +202FW | KRP2 +440RE | 238 |
| | mutant | KRP2 +202FW | LBb1.3 | 393 |

**Table S7**. Sequences for primers used for real-time RT-PCR.

| Gene | Primer name | Sequence |
|---|---|---|
| *BP* | BP-F | 5'-TGTTGTTTCCACATATGAGCTCTCT-3' |
| | BP-R | 5'-TCATGATCAGATCGGAAGCAAT-3' |
| *KNAT2* | KNAT2-F | 5'-TTCCGCTCGACGGAAGAC-3' |
| | KNAT2-R | 5'-AATCGGACGGCATCATCAAC-3' |
| *KNAT6* | KNAT6-F | 5'-GATGTCACCGGAGAGTCTCATG-3' |
| | KNAT6-R | 5'-CGGCGGAGGAACATAGCA-3' |
| *STM* | STM-F | 5'-CTCCTCCCCAAGGAACTAAGAAC-3' |
| | STM-R | 5'-TCCTCCTGCAACGATTTCG-3' |
| *KAN1* | KAN1-F | 5'-CCACGCGCGGTTTGTT-3' |
| | KAN1-R | 5'-CGACTTTGGAGTTGCTCTTTCA-3' |
| *KAN2* | KAN2-F | 5'-AAGGAACTAGATGGAAAGTGCTCAA-3' |
| | KAN2-R | 5'-GCTTGTTCCCGAGATGCTTG-3' |
| *FIL* | FIL-F | 5'-AAACCAACATGCCCCAACAG-3' |
| | FIL-R | 5'-TCACACCAACGTTAGCAGCTG-3' |
| *YAB5* | YAB5-F | 5'-ACGCCCTAATTTCCAGGCAAC-3' |
| | YAB5-R | 5'-GTTGCTCAGTTATGGTACGAG-3' |
| *ETT* | ETT-F | 5'-CGCCTACTCAATAACCGATCATC-3' |
| | ETT-R | 5'-ACGGCCCACACCAAATGTT-3' |
| *ARF4* | ARF4-F | 5'-CGCTTAAATCATTCCCGCAAT-3' |
| | ARF4-R | 5'-ACTTGTTGGCTTGGTAAGCAAAG-3' |
| *PHB* | PHB-F | 5'-GCTGTTGACTGGGTTCAGATGA-3' |
| | PHB-R | 5'-GCGAAATAGCGACTATGCCAAT-3' |
| *PHV* | PHV-F | 5'-GGCGGAGTTCCTTTGCAA-3' |
| | PHV-R | 5'-CCAGGCTTCATCCCAATCAT-3' |
| *ACT2* | ACT2-F | 5'-TCGGTGGTTCCATTCTTGCT-3' |
| | ACT2-R | 5'-GCTTTTTAAGCCTTTGATCTTGAGAG-3' |
| *EF1α* | EF1α-F | 5'-TGAGCACGCTCTTCTTGCTTTCA -3' |
| | EF1α-R | 5'-GGTGGTGGCATCCATCTTGTTACA -3' |
| *KRP5* | KRP5-F | 5'-TCCTAGTGTCAATCAATGTCAAACGGCAAAGT-3' |
| | KRP5-R | 5'-CGTCGTATCCGGCTCTAATTTCGAAATTAGAGCCGGA-3' |
| *KRP2* | KRP2-F | 5'-CGTGGATTTACGATGATTTGAA-3' |
| | KRP2-R | 5'-GCGGCGAGACTCTACATCTT-3' |
| *IPT3* | IPT3-F | 5'-CCGCCTGAAGCCGACTTAA-3' |
| | IPT3-R | 5'-TTTAGGACGGATTCAATGGAGAGA-3' |
| *KNOLLE* | KNOLLE-Q3 | 5'-TGCAGTCTTCAGCTCATTAGCTC-3' |
| | KNOLLE-Q5 | 5'-TGATGGTTGAATCGCAAGGTGAAC-3' |
| *HINKEL* | HINKEL_F | 5'-CCAAGCAGCGCATCCAA-3' |
| | HINKEL_R | 5'-AAGACTTGCCTAGAAGCTGAAAGC-3' |
| *histone H4* | histone H4-Q3 | 5'-TCCTTGAATGTTATCCCTCAGAAC-3' |
| | histone H4-Q5 | 5'-ACAGTCACAGATCTTTGGGTATC-3' |
| *CYCA3;1* | CycA3;1-Q3 | 5'-AGAAGTTTGTATTCTTCAGCAAC-3' |
| | CycA3;1-Q5 | 5'-TCTTCGTCAATTAGAGGTAAAGTC-3' |
